# Nucleus confinement within concave microcavities modulates nuclear morphology, subnuclear dynamics and mechanotransduction in human osteosarcoma cells

**DOI:** 10.64898/2026.03.20.712604

**Authors:** Ismail Tahmaz, Fabricio Frizera Borghi, Jean-Louis Milan, Philippe Kunemann, Tatiana Petithory, Mohammed Bendimerad, Valeriy Luchnikov, Karine Anselme, Laurent Pieuchot

## Abstract

Cells dynamically integrate biochemical and mechanical signals arising from their surrounding microenvironment to regulate morphology and behavior. Mechanical cues like matrix stiffness, surface topography, and other physical perturbations modify biophysical signals. Surface topography, particularly curvature regime acts as any important mediator of mechanotransduction by coordinating cytoskeletal organization, focal adhesion dynamics, and nuclear architecture. Curvature response has been demonstrated at broader length scales and influences nucleus shape change, chromatin organization, and gene regulation, positioning the nucleus as an active mechanosensitive hub. Bone tissue consists of a curvature-rich microenvironment defined by a trabecular architecture at tissue scale and by resorption cavities such as Howship’s lacunae at cellular scale. While these geometries are essential for homeostasis, their role in pathological context remains poorly understood. Osteosarcoma develops within this mechanically complex multiscale architecture, but how bone-inspired curvature regulates nuclear behavior and signaling in osteosarcoma cells remains unclear. Here, we engineered three-dimensional (3D) concave hemispherical substrates that recapitulate nucleus-scale bone micro-curvature and assessed their effects on human SaOS-2 osteosarcoma cells. In comparison with flat surfaces, concave confinement resulted in pronounced nuclear rounding and softening, accompanied by Lamin A/C reorganization and increased heterochromatin compaction marked by H3K9me3. Curvature-driven nuclear remodeling selectively modulated Hippo pathway main effectors YAP/TAZ without activating NF-κB mediated canonical inflammatory responses. Furthermore, cells maintained overall viability without elevated pathological DNA damage or apoptotic signaling, suggesting an adaptive, damage-tolerant nuclear response. Overall, these findings indicate nucleus-scale curvature as a critical regulator within the bone microenvironment that governs nuclear modelling and mechanosensitive signaling in osteosarcoma cells. Incorporating physiologically relevant geometry into *in vitro* models establishes new insight into cancer microenvironment crosstalk and highlights nuclear interior and outer architecture as a key regulator of tumor cell behavior.

## 1. Introduction

The cellular and tissue specific microenvironment provides a dynamic combination of signals tightly regulating cell behavior and morphology. Although biochemical cues such as extracellular matrix composition, growth factors, and cytokines—play a critical role, their effects are closely intertwined with the physical signals from the surrounding tissue ^(1,2)^. Mechanical cues like matrix stiffness, surface topography, and other physical perturbations activate mechanotransduction pathways, which in turn influence cell polarization, migration and fate decision ^(3–5)^. Apart from other mechanical factors, regarding cell-material interaction, topographic cues serve as an important geometric cue influencing cytoskeleton, transmitting physical information to the nucleus either directly or indirectly, thus linking extracellular geometry to nuclear architecture and gene regulation ^(6,7)^. While surface topography includes a wide range of geometrical features, curvature serves a powerful and biologically relevant topographic cue, as it naturally creates geometric confinement and guides the mechanical forces experienced by cells.

Curvature sensing can occur across multiple length scales. Curvature at both the cellular and subcellular levels can guide cells to sense mechanical cues. It reorganizes focal adhesion dynamics, cytoskeleton, and nuclear morphology, in turn affecting fundamental cellular behaviors such as migration, differentiation, and fate decisions ^(8,9)^. Earlier studies investigating curvature landscapes both *in vitro* and *in silico*, combining Gaussian curvature—convex and concave, and microgrooves have shown that cell-scale curved geometry reorganizes the cytoskeleton and focal adhesions, resulting in pronounced nuclear elongation or compression depending on the local geometry of the substrate ^(10–13)^ . Such geometric cues not only change cell morphology but also direct migration trajectories and osteogenic differentiation through nucleus-cytoskeleton coupling ^(11,12,14)^ . It was shown that cells prefer migrating towards concave surfaces, where they spread radially, while convex surfaces promote osteogenesis by compressing the nucleus. *Vassaux et al.* (*2019*) and *Pieuchot et al.* (*2018*) combining in silico and in vitro approaches further demonstrated that nuclear stiffness and its mechanical coupling with other subcellular structures serves as an additional crucial determinant of curvature readout, alongside the contribution of the cytoskeleton ^(11,15)^. Furthermore, *Leclech et al.* (*2025*) recently revealed that surface adhesion forces contribute more to nuclear deformation within microgrooves than the cytoskeleton does ^(16)^. Collectively, although curvature sensing seems to be a highly intricate process involving both the cytoskeleton and focal adhesion proteins, the nucleus plays equally important role receiving transmitted physical signals from cytoskeleton. It responds to mechanical cues and its shape can be accompanied by changes in chromatin organization, gene regulation, and cell cycle progression for longer term.

Since the surrounding topography significantly affects force distribution inside the cell nucleus, changes in nuclear morphology driven by curvature remodel lamin organization and chromatin compaction ^(12,17)^. These interactions mediate mechanotransductive signalling and recruit Lamin A/C which are integral mediators of the nuclear envelope. Lamin A/C responds to substrate curvature by tuning nuclear stiffness and cellular tension ^(18–21)^. Besides, changes in nuclear morphology have been associated with heterochromatin and histone marks remodeling such as H3K9me3, and accessibility of transcription factors ^(22–24)^. Hence, excessive or uneven nuclear stress might result in laminopathic nuclear morphologies, cause DNA damage, and trigger stress-response signaling pathways ^(25–27)^.

Beyond cell and nuclear scale, interest in mesoscale curvature has grown rapidly in recent years to characterize many native tissues. Previous studies have shown that cells are remarkably sensitive to their mesoscale environment. The shape of these geometries can coordinate collective cell behavior, control how cells are spatially organized within a scaffold, and influence their proliferation ^(28–31)^. Numerous studies have been investigating how curvature affects biological structures such as blood vessels, trabecular bone, and early embryonic formations like the blastocyst. For example, varying curvature degree present in blood vessels leads to non-uniform shear stress, stimulating gene expression that promotes vascular remodeling and proliferation ^(32–34)^. Another notable example also revealed that curvature plays also a crucial role in developmental processes such as embryogenesis. A recent study designed and developed a surface mimicking the *in vivo* geometry of blastocysts, consisting of stochastically distributed concavities ^(35)^. This surface was able to reprogram primed stem cells back into a naïve state in vitro by inducing apical constriction, strengthening E-cadherin/RAC1 signaling, activating YAP-dependent histone modifications, and increasing naïve state marker–NANOG expression which in turn represent the higher developmental potential and stemness. Lastly, the 3D-printed scaffold, inspired by the hyperboloidal topography of bone and characterized by negative Gaussian curvature, enhanced mesenchymal stem cell differentiation toward osteogenesis ^(36)^. This finding exerts the importance of the cellular microenvironment in regulating bone formation and remodeling. Indeed, bone remodeling depends on the homeostasis coordination among osteoclasts, osteocytes and osteoblasts ^(37)^. Briefly, osteoclasts create the concave pits i.e–Howship’s lacunae during active bone resorption. Osteoblasts primarily sense the resulting pit curvature/concavity afterward as mechanical cues. After that, they respond by filling in these pits during the regeneration phase. The effects associated with this specific curvature sign have been the focus of considerable study and modeling efforts in recent years, especially regarding the mechanobiological responses of preosteoblasts on concave geometries in the context of tissue regeneration ^(28,38,39)^. While these cues are essential for maintaining homeostasis, it should be also considered in pathophysiology. Cancer cells respond to these signals and leveraged them to support initiation or progression of malignant states ^(40)^. Thus, engineered substrates recapitulate the importance of physiological curvature and extend its relevance from tissue engineering to disease context, including understanding crosstalk between cancer and its physical microenvironment.

Osteosarcoma is a highly aggressive primary bone malignancy that arises within a mechanically complex and topographically rich microenvironment ^(41)^. Native bone is defined by trabecular curvature and resorption cavities such as Howship’s lacunae, which present locally either concave or saddle-like geometries at the cellular scale ^(42–44)^. Osteogenic and malignant cells experience spatial confinement and altered mechanical loading in such bone-niche geometry. Seminal study has shown that the geometry of the surrounding environment where osteosarcoma cells are constricted strongly influences how forces are distributed within the cell nucleus. This process depends on actomyosin contractility, vimentin, and LINC complexes, which together play a crucial role in driving nuclear deformation of metastatic osteosarcoma cells during interactions with micropillars ^(45,46)^. Together, these mechanisms position nuclear mechanics as a central role in regulating osteosarcoma cell behavior in response to their physical microenvironment.

Irregular nuclear shapes are a hallmark of cancer and often considered poor prognosis ^(47–49)^. Despite extensive investigation into biochemical drivers of osteosarcoma progression, the role of bone-inspired curvature and mechanical confinement in regulating osteosarcoma cell behavior remains still unclear. Since this microenvironment presents distinct geometric cues that are difficult to reproduce using conventional two-dimensional (2D) cell culture systems, we deployed maskless grayscale lithography (GSL) enabling rapid prototyping and 3D structure ^(50)^. To systematically explore how surface geometry affects nuclear shape and related subnuclear processes, we used SaOS-2 human osteosarcoma cells, which are commonly employed in bone tumor mechanobiology studies due to their well-characterized nuclear deformability in response to mechanical cues ^(45,46,51)^. The eukaryotic cell nucleus is typically reported to be about 6 µm in diameter ^(52)^, which would allow it to be readily accommodated within a cell-scale bone microenvironment. By considering both the physiological properties of osteosarcoma nuclei and the curvature of Howship’s lacunae ^(53,54)^, we designed a 3D micro hemisphere surface that consists of negative Gaussian/concave curvature. This approach allowed us to explore how cell-scale bone mimetic surfaces could be designed to guide specific subcellular behaviors, and might lead to new directions in cancer research.

In this study, we engineered 3D array of concave hemispheres to mimic local cell nucleus microenvironment in Howship’s lacunae and to investigate how curvature affects SaOS-2 osteosarcoma cells subnuclear dynamics compared with flat substrates. We analyzed nuclear morphology; lamina and epigenetic remodeling (Lamin A/C, H3K9me3); mechanotransduction and inflammatory pathways (YAP/TAZ and NF-κB); DNA damage (γH2AX); and viability along with apoptotic signaling (cleaved caspase-3). Our results reveal that concave confinement resulting in nuclear rounding induces lamina softening and promotes distinct modulation of Hippo pathway components (YAP/TAZ) without NF-κB activation. This adaptive response also enhances chromatin compaction rather than global deacetylation and damage-tolerant response. Taken together, these findings indicate that nucleus scale curvature acts as a crucial signal within the bone microenvironment, influencing how osteosarcoma cells organize their nuclei in their physical microenvironment. This allows us to better understand the crosstalk among the cancer microenvironment, nuclear shape, and cell behavior.

## 2. Results

### 2.1. The predetermined surface parameters particularly lateral diameters are successfully achieved

To mimic cell nucleus-scale curvature of Howship’s lacunae, an array of concave lodges was therefore generated using maskless GSL in varying height and width ranges to represent physiologically relevant scales (Fig. 1A). Howship’s lacunae in human trabecular bone exhibit mean maximum depths of approximately 30 µm and widths of several hundred micrometers as estimated from reported surface areas ^(53,54)^, and here we engineered a series of hemispherical concave surfaces whose depths ranged from 5, 7.5, 10 and 15 microns and widths twice each, respectively (Fig. 1B-C). The use of maskless greyscale lithography allowed rapid prototyping of hemisphere patterns representing down-scaled and nucleus scale analogues of this resorptive geometry. After intended surfaces were designed, the final surface topography was characterized using scanning electron microscopy (SEM) and profilometry to assess fidelity between design and fabricated structures. SEM imaging revealed well-defined, homogeneously distributed concave hemispherical surfaces across all designed groups. The hemispheres exhibited smooth curvature and clear boundary definition, with minimal collapse or deformation (Fig. 1D). After that, surface profiles were obtained by profilometer with cross-sectional readout to quantify resulting depth and diameter (Fig. 1E). The difference between designed and resulting diameter was minimal across all groups, confirming highly achieved dimensional fidelity between fabrication and design.

**Figure 1.**
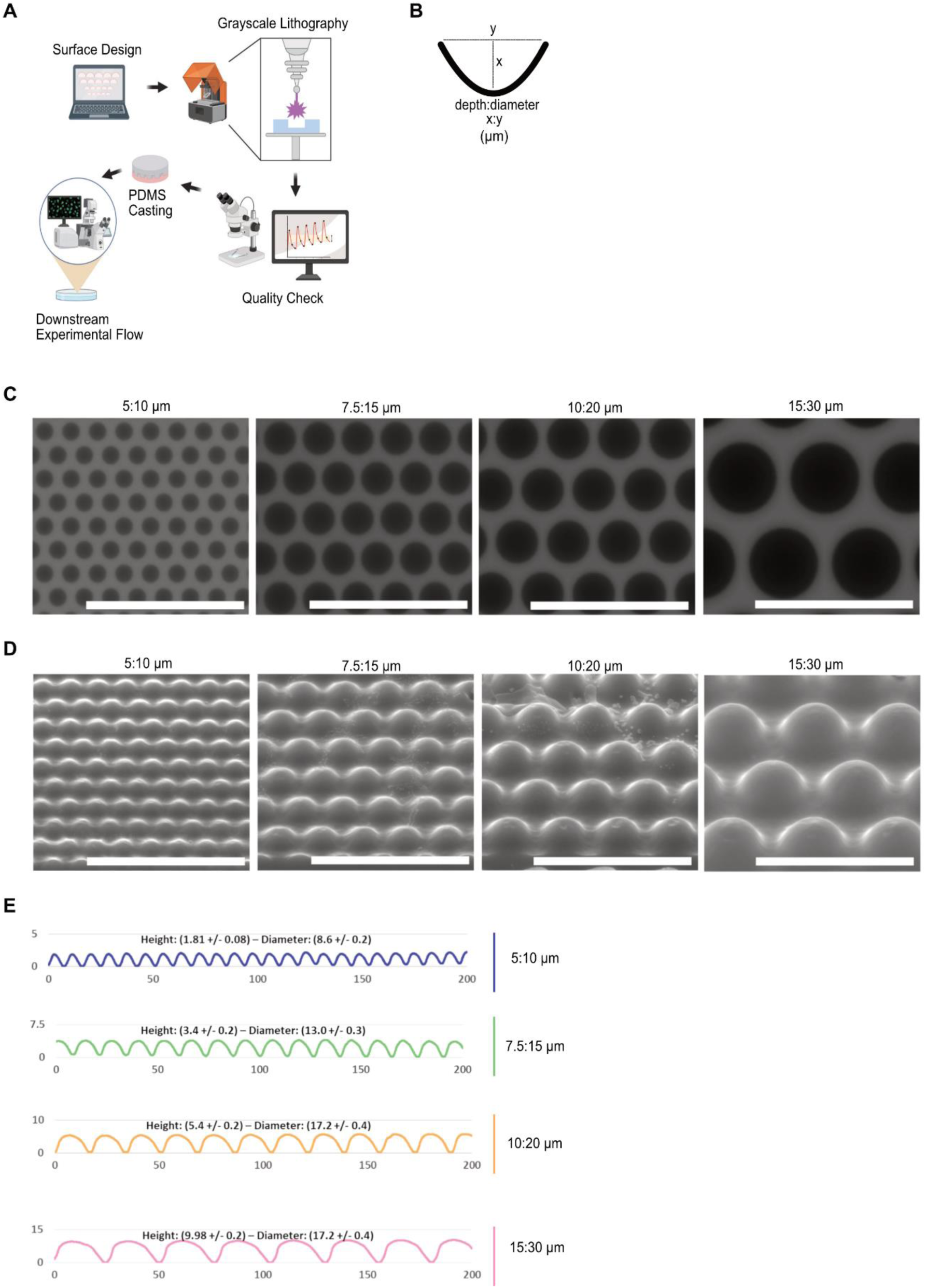
The fidelity between designed and fabricated structures is largely achieved **(A)** Schematic illustration of the overall workflow from the initial design to the fabrication of hemispherical surfaces. **(B)** Definition of hemisphere surface diameter and depth used in this study. **(C)** Digital mask and **(D)** SEM images of varying length of hemisphere surfaces. Scale bar, 50 µm. **(E)** Surface profiles of varying length of hemisphere surfaces.

These results confirm that the fabricated surfaces provide reproducible and well-defined concave hemisphere geometries with largely preserved lateral and planar dimensions.

### 2.2. Substrate local curvature influences nucleus sphericity

Following the design and fabrication of the biomimetic surfaces, we investigated how hemisphere diameter and depth specifically influence nuclear geometry. To this end, we fabricated hemispherical patterns with varying dimensions, with depths ranging from 5 to 15 µm and corresponding diameters from 10 to 30 µm. Nuclear sphericity was assessed by both (2D) and 3D analyses, including consideration of nuclear area, volume, and aspect ratio. The 3D analysis showed that nuclei located within larger hemispherical patterns (10 and 15 µm diameters) occupied notably smaller nuclear area and volume compared with nuclei on smaller hemispheres and flat surfaces (Fig. 2A; *see SI*, Fig. S1A-B). Consequently, nuclei experiencing these larger hemispherical curvatures exhibited a higher 3D nuclear sphericity index, indicating a strong relationship between pattern diameter and nuclear morphology (Fig. 2B). These 3D findings were also consistent with the 2D analyses of nuclear area and sphericity (*see SI*, Fig. S1C). In the 2D analysis, larger hemispherical diameters promoted less nuclear spreading and significantly higher nuclear circularity (Fig. 2C; *see SI*, Fig. S1C-D). Among the larger patterns, nuclei confined within 10 µm hemispheres exhibited greater roundness, characterized by a more evenly distributed aspect ratio, compared with those within 15 µm hemispheres (*see SI*, Fig. S1E). Notably, in the initial surface design, many nuclei were not positioned within hemispherical features. However, cells on 10 µm diameter patterns showed a higher propensity for nuclei to reside within the hemispheres than those on 15 µm patterns (Fig. 2D). We chose hemispherical patterns with a height of 10 µm and a width of 20 µm for further experiments based on these findings.

**Figure 2.**
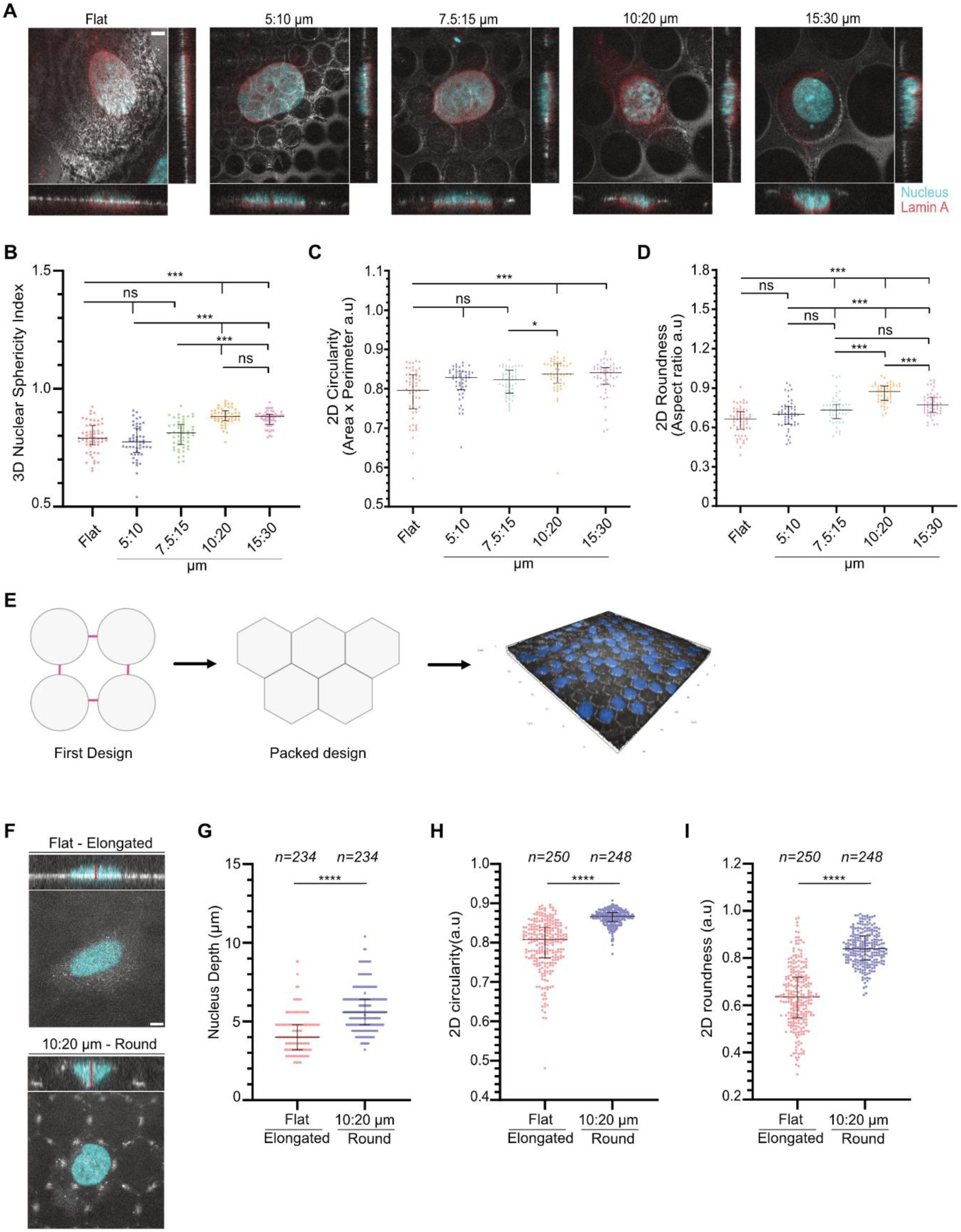
Nuclei on larger hemispherical substrates exhibited higher sphericity and roundness, particularly within the 10 µm depth and 20 µm diameter pattern. **(A)** Representative immunofluorescence images of nuclei (Hoeschst, light blue), Lamin A (red) of SaOS-2 cells cultured on flat PDMS surfaces (elongated nuclei), and varying length of concave hemispherical PDMS surfaces. Corresponding DIC images are shown. Scale bar, 5 µm. **(B)** Quantification of 3D nucleus sphericity based on volume and perimeter, **(C)** 2D circularity based on area and perimeter and, **(D)** 2D nucleus roundness based on aspect ratio revealed that larger diameter surfaces especially 10 µm depth, 20 µm diameter resulted in uniform and rounder nuclei morphology. **(A-D)** Experiments were performed only once. **(E)** Illustration of design optimization: reducing flat regions between hemispheres in the 10:20 µm layout improved uniform nuclear distribution across the reconstructed surface. **(F)** Representative orthogonal view of nuclei (Hoeschst, light blue) of SaOS-2 cells cultured on flat PDMS surfaces (elongated nuclei), and 10:20 µm hemisphere PDMS surfaces with corresponding DIC images. Scale bar, 5 µm. **(G)** Quantification of 3D nuclei depth, **(H)** 2D circularity and **(I)** 2D roundness measurement confirm that nuclei enclosed within 10:20 µm packed hemispherical design giving higher 3D depth maintained greater nuclear circularity and roundness. **(F-I)** “n” represents the number of nuclei analyzed from six independent experiments. (*p < 0.05; **p < 0.01; ***p < 0.001; ****p < 0.0001; n.s., not significant).

By reducing the flat areas between hemispheres, this optimized design improved uniform mechanical confinement (Fig. 2E). The resulting geometry likely imposed more uniform nuclei confinement within surfaces, promoting a rounder nuclear shape. SaOS-2 cells grown on within concave hemispherical polydimethylsiloxane (PDMS) surfaces (10 µm depth, 20 µm in diameter) exhibited 3D nuclear confinement that was apparent in both orthogonal and reconstructed views (Fig. 2F). In these conditions, the nuclei exhibited nearly full confinement within the hemispherical concavities, adopting a rounded shape that closely conformed to the surface curvature. The nuclear depth nearly matched the designed 10 µm feature depth, and both nuclear sphericity and roundness were consistently and significantly higher than in cells on flat PDMS, reflecting a more uniform morphology (Fig. 2G–I). By contrast, cells on flat fibronectin-coated PDMS spread along the planar surface, with nuclei extending mostly laterally and showing minimal vertical dimension. As a result, nuclear depth, sphericity, and roundness were all considerably lower in the flat condition compared to nuclei confined in the hemispheres. This striking difference in nuclear geometry highlights the 3D nature of the confinement, suggesting that all subsequent analyses are strongly shaped by the spatial configuration of the nucleus.

Overall, our findings showed a strong relationship between hemispherical geometry and nuclear morphology by consistently showing that nuclei that enclosed within hemispherical surfaces exhibit increased nuclear sphericity and roundness. Furthermore, nuclei on hemispherical surfaces showed higher nuclear depth than nuclei on flat substrates, suggesting a 3D elastic response of the nucleus confinement within the hemispherical geometry.

### 2.3. The concave surface promotes transition to a softer nuclear state in osteosarcoma cells

Next, we analyzed nuclear integrity in response to concave curvature. We therefore focused our analysis on the mechanosensitive perinuclear lamin proteins—specifically lamin A and lamin C which maintain nuclear architecture and structural integrity. To compare potential differences between round and elongated nuclei, we performed immunostaining for lamin A. Our results indicated that lamin A levels did not differ significantly between these two nuclear shapes (Fig. 3A-B). In addition to lamin A, we examined the other major isoform, lamin C, alongside lamin A. Interestingly, unlike lamin A alone, the combined involvement of lamin A and lamin C was markedly higher in nuclei cultured on flat surfaces. This observation suggests that nuclei on hemispherical surfaces do not need to maintain mechanical stability and integrity, leading mechanically softer than those on flat surfaces (Fig. 3C-D). Moreover, our results suggest that nuclear shape changes induced by confinement within concave structures differentially regulate Lamin A and Lamin C levels and may not directly correlate with Lamin A alone.

**Figure 3.**
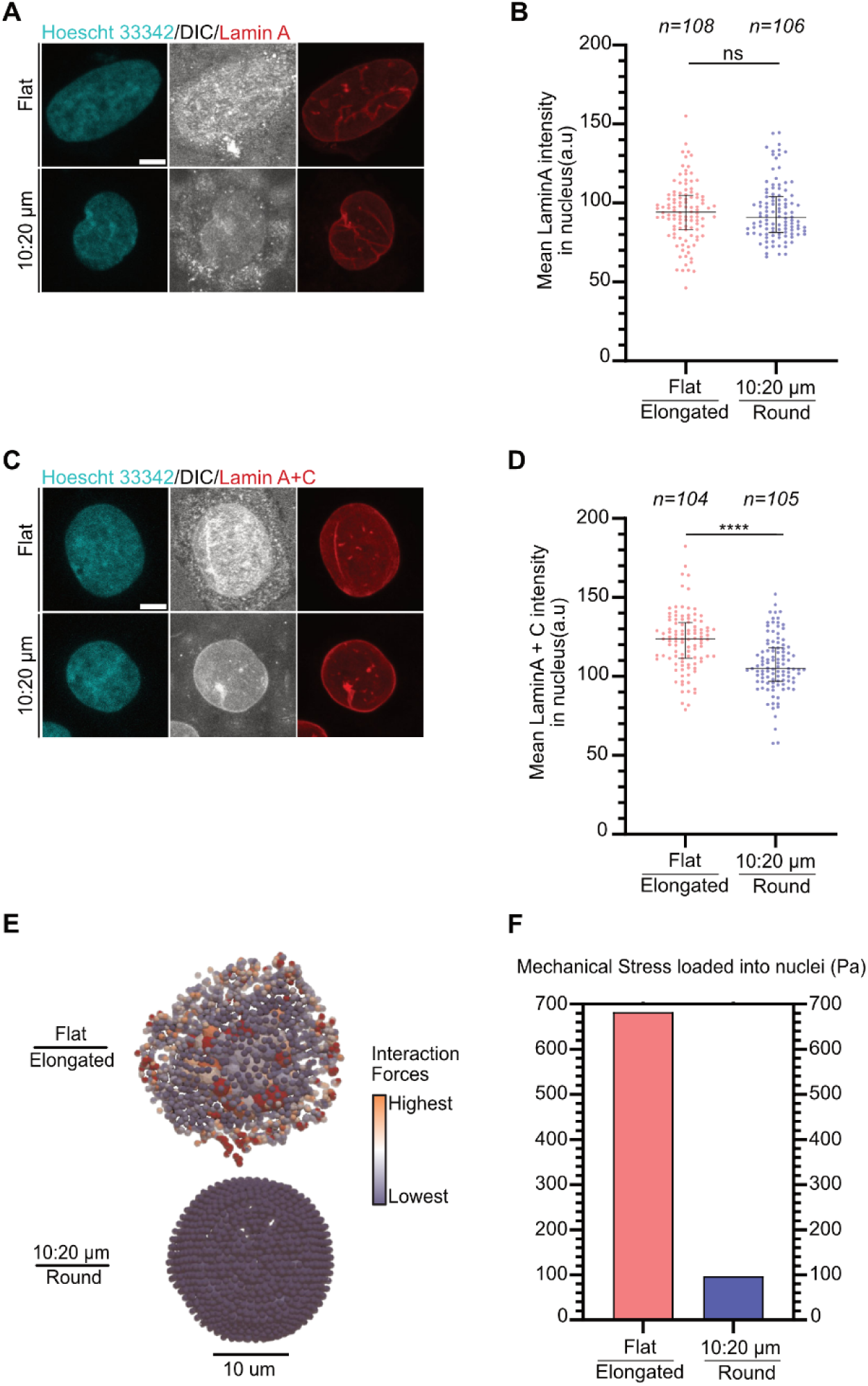
Nuclear rounding within hemisphere surfaces perturb Lamin A/C but not Lamin A levels and redistribute mechanical stress in SaOS-2 cells. **(A, C)** Representative fluorescence images of SaOS-2 cells cultured on fibronectin-coated flat PDMS surfaces (elongated nuclei) or concave hemispherical PDMS substrates (10 µm depth, 20 µm diameter; round nuclei) stained for **(A)** Lamin A (red) or **(C)** Lamin A/C (red) and Hoechst (light blue); corresponding DIC images are shown. **(A, C)**. Scale bar, 5 µm. **(B, D)** Quantification of mean nuclear fluorescence intensity revealed no significant difference in Lamin A levels between flat and hemispherical surface **(B)**, whereas Lamin A/C levels were significantly reduced in mechanically confined, round nuclei **(D)**. **(B, D)** “n” represents the number of nuclei analyzed from three independent experiments. (****p < 0.0001; n.s., not significant). **(E)** Stress simulation maps showing bead–nucleus interaction forces in elongated (flat) and round (hemispherical) nuclei. Scale bar, 10 µm. **(F)** Quantification of simulated mechanical stress revealed higher stress accumulation in elongated nuclei compared with round nuclei, indicating reduced nuclear stress under hemispherical confinement.

We further performed finite element modeling to simulate the stress distribution along the elongated and round nuclei on flat and hemispherical surfaces, respectively (Fig. 3E). The computational analysis revealed that mechanical stress was profoundly higher in elongated nuclei on flat surfaces, whereas round nuclei within hemispherical surfaces exhibited much lower stress accumulation as a result of more homogeneous stress distribution (Fig. 3F). This observation indicates that elongated nuclei on flat surfaces exhibit higher mechanical rigidity, consistent with their elevated Lamin A/C levels. In contrast, round nuclei confined within concave surfaces undergoes mechanosensitive adaptation enabling a softer and likely more compliant nuclear state with lower Lamin A/C abundance.

### 2.4. Nuclear rounding increases chromatin condensation through repressive heterochromatin formation

To investigate how nuclear confinement in hemispherical patterns affects the chromatin regulatory landscape and associated epigenetic modifications, we first examined key repressive epigenetic markers. We tried to achieve integrated view of the epigenetic state by assessing histone deacetylase 1 (HDAC1) activity—which removes acetyl groups from lysine residues on histone tails—and by analyzing the repressive mark H3K9me3, representing trimethylation at lysine 9 of histone H3 ^(55,56)^. Both H3K9me3 and HDAC1 are established mediators of transcriptional repression. We quantified the mean fluorescence intensity (MFI) of both markers in nuclei exhibiting round and elongated morphologies. To begin with, HDAC1 fluorescence intensities were comparable between conditions, indicating a similar global deacetylation profile in both nuclear shapes (Fig. 4A). Consistently, quantitative image analysis showed no significant difference in average HDAC1 levels between nuclei grown on flat and hemispherical substrates (Fig. 4B). Unlike HDAC1, H3K9me3 signals were visibly elevated in round nuclei enclosed within the hemisphere surface (Fig. 4C-D). Increased H3K9me3 levels reveal that chromatin compaction under mechanical constraints promotes the formation of repressive heterochromatin.

**Figure 4.**
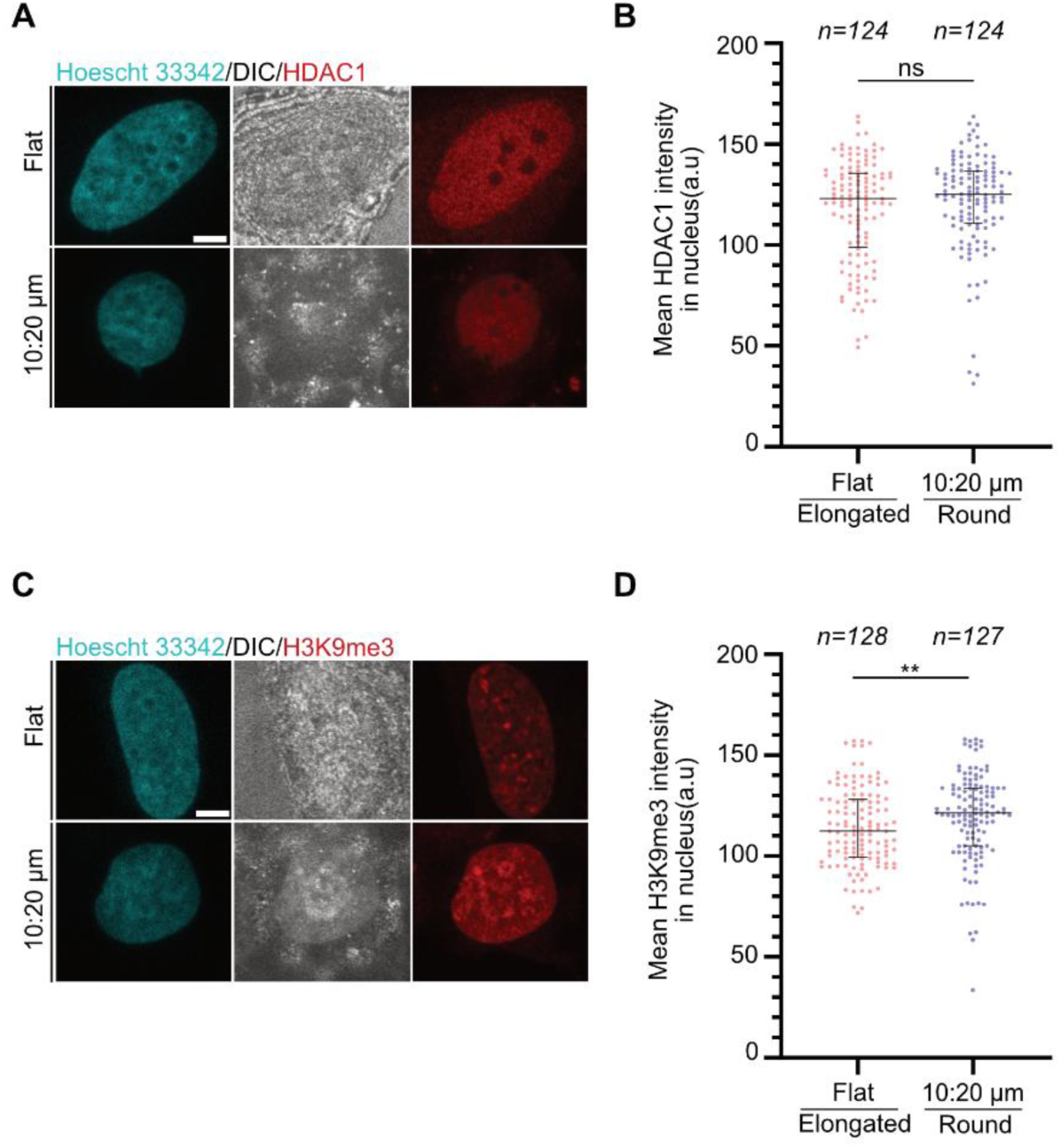
Nuclear rounding within hemisphere surfaces selectively regulates repressive epigenetic markers. **(A, C)** Representative fluorescence images of SaOS-2 cells cultured on fibronectin-coated flat PDMS surfaces (elongated nuclei) or concave hemispherical PDMS substrates (10 µm depth, 20 µm diameter; round nuclei) stained for **(A)** HDAC1 (red) or **(C)** H3K9me3 (red) and Hoechst (light blue) for nucleus with corresponding DIC images. **(A, C)**. Scale bar, 5 µm. **(B, D)** Quantification of mean nuclear fluorescence intensity revealed no significant difference in HDAC1 levels between flat and hemispherical surfaces **(B)**, whereas H3K9me3 levels were significantly increased in mechanically confined, round nuclei **(D)**. **(B, D)** “n” represents number of nuclei analyzed from three independent experiments. (**p < 0.01; n.s., not significant).

Taken together, these findings revealed that mechanical confinement resulted in rounder nuclei changes and promotes selective enhancement of repressive heterochromatin formation through perinuclear lamina–chromatin interactions, rather than via HDAC1-mediated deacetylation.

### 2.5. Concave local curvature induces sublethal DNA damage with preserved viability

Following epigenetic remodeling, we next assessed DNA damage and functional outcomes of nuclear confinement. We evaluated mean nuclear γH2AX intensity as the primary readout rather than foci, since UV irradiation resulted in γH2AX foci fusion (Fig. 5A; *see SI*, Fig. S3). The mean nuclear γH2AX fluorescence intensity was slightly higher in round nuclei within concave surface compared to elongated nuclei on flat PDMS (Fig. 5B). In both conditions, γH2AX levels were much lower than the irradiated control samples, suggesting that the cells are undergoing only mild DNA stress without developing significant genomic instability within the hemisphere surface.

**Figure 5.**
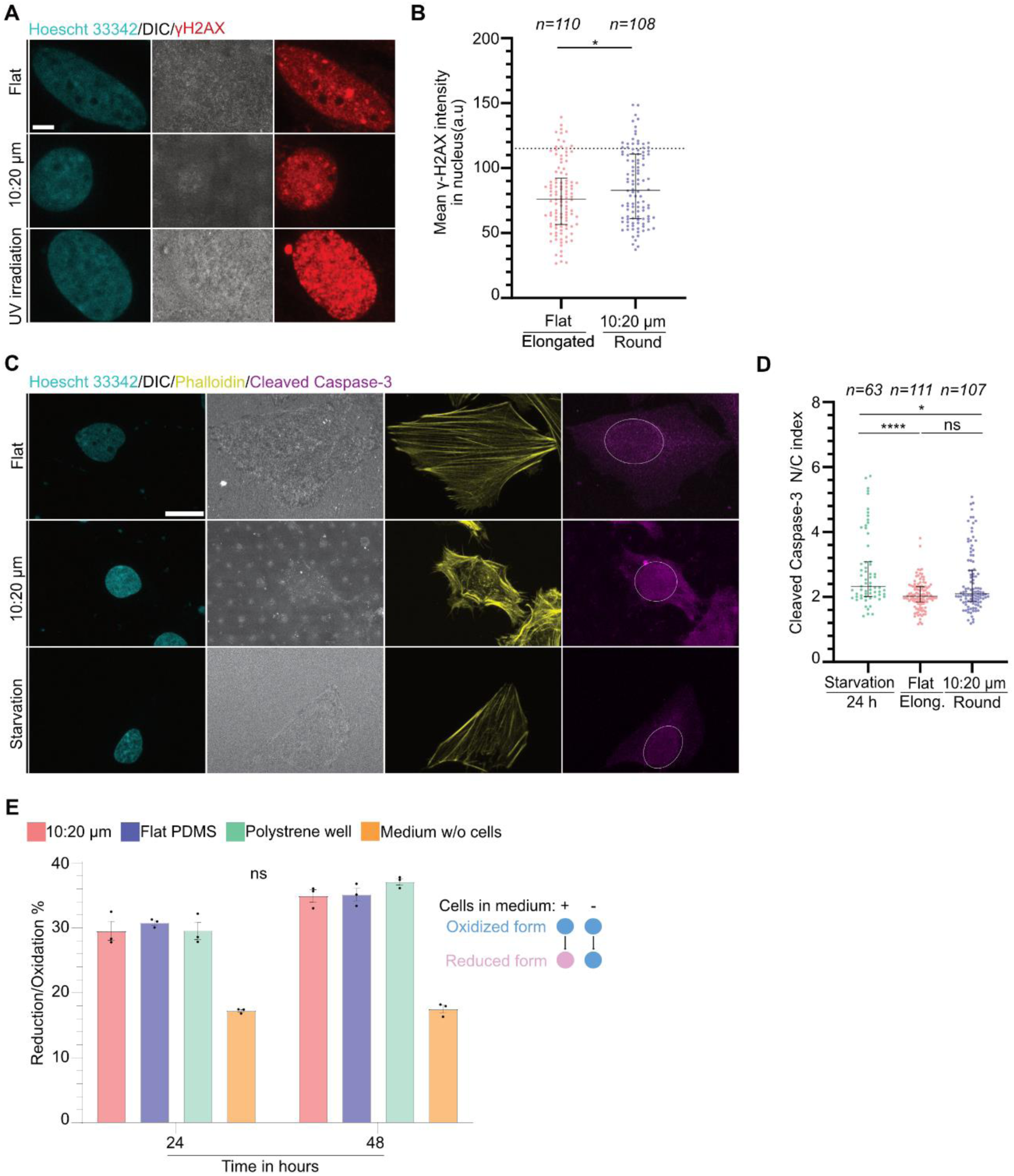
Nuclear rounding causes sublethal DNA damage in response to concave geometry without affecting survival in SaOS-2 cells. **(A)** Representative fluorescence images of SaOS-2 cells cultured on flat PDMS surfaces (elongated nuclei), concave hemispherical PDMS substrates (10 µm depth, 20 µm diameter; round nuclei) and irradiated sample stained for γH2AX (red) and Hoechst (light blue) for nucleus; corresponding DIC images are shown. Scale bar, 5 µm. **(B)** Quantification of γH2AX (MFI) revealed a slight but significant increase in mechanically confined round nuclei compared with flat counterparts, while remaining lower than irradiation-treated positive controls, indicating modest DNA damage under nuclear confinement rather than pathological damage. The dashed line represents MFI of irradiated samples. **(C)** Representative immunofluorescence images of cleaved caspase-3 in SaOS-2 cells cultured on flat, concave hemispherical PDMS surfaces and cells with serum and nutrient starvation (24 h) included as an apoptosis-inducing positive control. Samples were stained for cleaved caspase 3 (purple), Hoechst (light blue) for nucleus and Phalloidin (yellow) for F-actin; corresponding DIC images are shown. Scale bar, 20 µm. **(D)** Nuclear-to-cytosol ratio of cleaved caspase-3 MFI reveals no significant difference between elongated and round nuclei; both condition exhibits significantly lower levels than starvation controls, indicating that nuclear confinement does not promote apoptotic commitment. **(B, D)** “n” represents the number of nuclei analyzed from three independent experiments. (*p < 0.05; (****p < 0.0001; n.s., not significant.) **(E)** Cell viability assessed by Alamar Blue assay showed no significant difference between cells cultured on flat and concave hemispherical PDMS surfaces. Reduction/oxidation ratios were comparable to those of cells cultured on polystyrene (positive control) and higher than cell-free medium, indicating preserved metabolic activity under nuclear confinement. Viability assay was performed three times. (n.s., not significant.)

DNA damage commonly induces apoptosis via p53-dependent signaling pathways and subsequent caspase activation ^(57)^. Accordingly, levels of cleaved caspase-3 which is the executioner caspase activated by downstream initiator caspases-8 and -9 were quantified to determine whether confinement-induced γH2AX reflects functionally lethal DNA damage or not ^(58)^. The nuclear-to-cytosolic ratio of cleaved caspase-3 is a reliable marker of apoptotic cleavage. As a physiological positive control, cells were subjected to 24 hours of serum and nutrient starvation, a condition known to activate both intrinsic (mitochondrial/Bak-mediated) and extrinsic (death receptor–mediated) apoptotic pathways in cancer cells ^(59)^. Despite elevated γH2AX levels, cells with elongated and round nuclei exhibited comparable nuclear-to-cytosolic cleaved caspase-3 ratios (Fig. 5C), indicating no difference in apoptotic effector localization (Fig. 5D). Although cleaved caspase-3 levels in both conditions were markedly lower than in cells exposed to starvation medium, flat controls displayed even lower levels compared with the starvation control, confirming minimal baseline apoptosis. Collectively, these results demonstrate that confinement enhances sublethal DNA damage signaling without promoting caspase-3 progression toward apoptosis.

We next evaluated cellular metabolic fitness to confirm the maintenance of cell survival. Alamar blue assay was employed to quantitatively assess cell viability based on subcellular redox reactions. SaOS-2 cells were treated with Alamar Blue reagent, and metabolic activity was evaluated by measuring absorbance. Cells cultured on polystyrene (control group) exhibited viability similar to tested experimental conditions and all conditions have given greater absorbance values than culture medium without cells confirming the absence of metabolic activity (Fig. 5E). In summary, these results revealed that SaOS-2 cells maintain viability on both hemispherical and flat PDMS surfaces, suggesting that the hemispherical surface did not adversely affect overall osteosarcoma cell viability.

Together, these data reveal that nuclear rounding as a result of geometric confinement within the biomimetic hemisphere surface enhances a protective adaptation resulting in sublethal DNA damage. without apoptotic triggering.

### 2.6. Concave hemisphere surface differentially regulates YAP/TAZ subcellular localization independently of canonical inflammatory signaling

Since nuclear dynamics are closely linked to the activity of key mechanosensitive transcription factors—Yes-associated protein (YAP) and transcriptional coactivator with PDZ-binding motif (TAZ), the principal effectors of the Hippo pathway ^(60)^—we first examined the active, dephosphorylated forms of YAP and TAZ during their nuclear translocation, a process highly sensitive to mechanical cues. In this experimental setup, soft PDMS substrates were used as control conditions, as commonly applied in YAP/TAZ studies, to enable comparison between nuclear and cytoplasmic localization. The elastic properties of these substrates were validated by atomic force microscopy (AFM), and the measured values closely matched previously reported data (*see SI*, Fig. S5) ^(61)^. Across all experimental conditions, both YAP and TAZ predominantly localized to the nucleus on soft PDMS substrates (Fig. 6A–D). In flat surface that induced elongated nuclear morphology, YAP localized predominantly to the nucleus, consistent with canonical stretch-induced activation (Fig. 6A-B). Strikingly, however, YAP was significantly retained in the cytoplasm in cells with round nuclei confined within concave hemispheres, indicating geometry-dependent cytoplasmic sequestration. Conversely, TAZ exhibited the opposite trend. Its nuclear accumulation was markedly enhanced in mechanically confined, round nuclei compared with flat controls (Fig. 6C-D). Quantification of nuclear-to-cytoplasmic mean intensity ratios confirmed this different trend of both mechanotransducers (Fig. 6A–D). These findings suggest that nuclear rounding induced by hemispherical confinement triggers a distinct mechanotransduction signature that differentially regulates YAP and TAZ.

**Figure 6.**
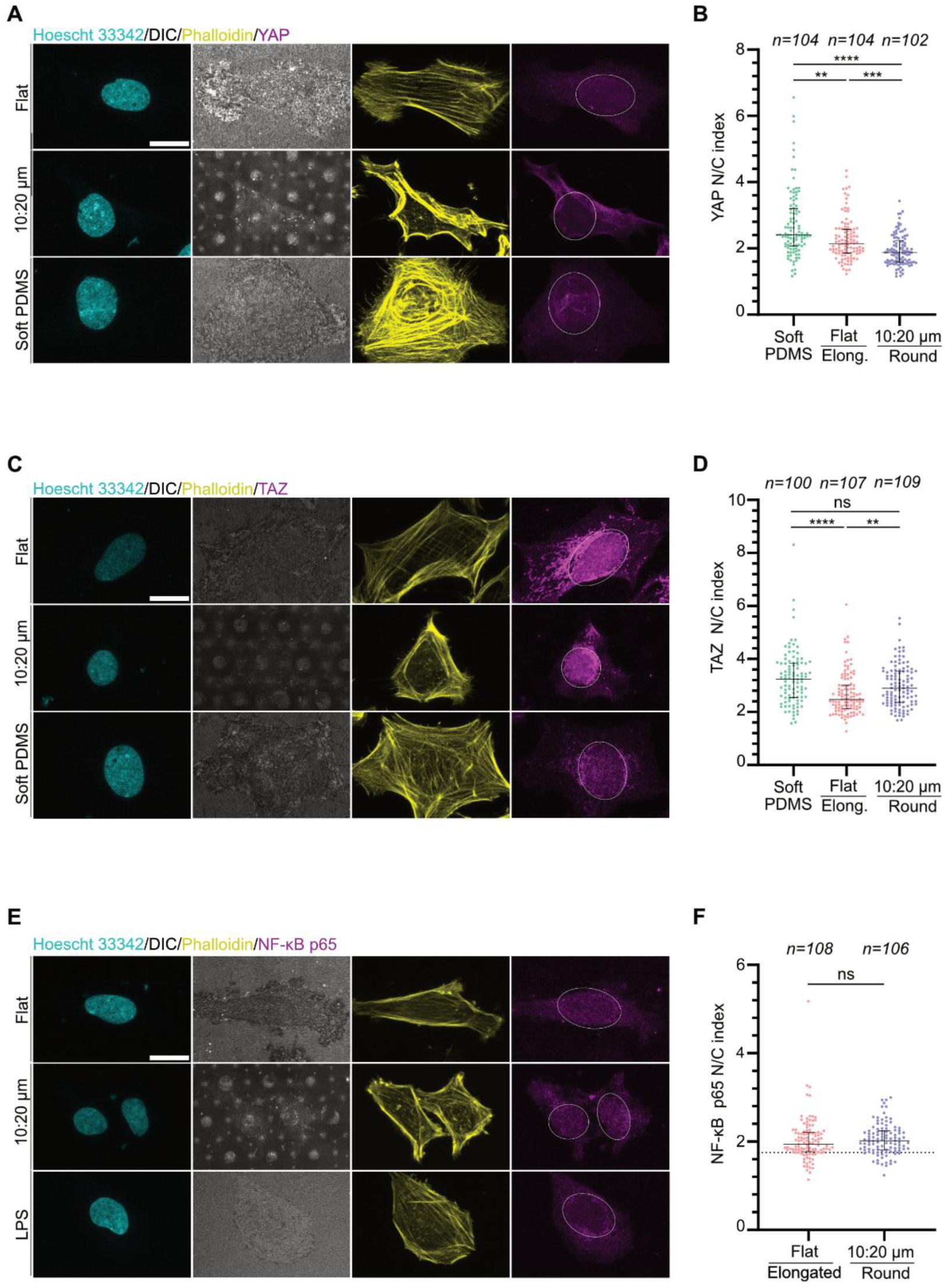
Nuclear shape–dependent mechanotransduction selectively modulates YAP/TAZ localization independently of NF-κB signaling. **(A, C)** Representative immunofluorescence images of YAP **(A)** and TAZ **(C)** in SaOS-2 cells cultured on flat PDMS surfaces (elongated nuclei), concave hemispherical PDMS surfaces (10 µm depth, 20 µm diameter; round nuclei) and cells on soft PDMS surfaces stained for YAP or TAZ (purple), Hoechst (light blue) for nucleus and Phalloidin (yellow) for F-actin; corresponding DIC images are shown. **(A, C)** Scale bar, 20 µm. **(B, D)** Quantification of nuclear-to-cytosolic MFI ratios for YAP **(B)** and TAZ **(D)**. YAP is significantly more cytosolic in round nuclei formed within concave hemispheres compared to elongated nuclei on flat PDMS. Cells on soft PDMS display significantly higher nuclear YAP localization than both elongated and round nuclear conditions. TAZ nuclear localization is significantly increased in round nuclei compared to elongated nuclei on flat PDMS. Soft PDMS substrates also exhibit high nuclear localization of TAZ. **(E)** Representative immunofluorescence images of NF-κB p65 in SaOS-2 cells cultured on flat PDMS, concave hemispherical PDMS surfaces and cells treated with LPS for an hour as a positive control stained for stained for NF-κB p65 (purple), Hoechst (light blue), and F-actin (Phalloidin, yellow). Corresponding DIC images are shown. Scale bar, 20 µm. **(F)** Quantification of NF-κB p65 nuclear-to-cytosolic MFI ratio. No significant difference in NF-κB p65 nuclear localization is observed between elongated and round nuclei. The dashed line represents MFI of samples treated with LPS whose NF-κB nuclear localization remains lower than that observed under both elongated and round nuclear conditions in osteosarcoma cells. **(B, D, E)** “n” represents the number of nuclei analyzed from three independent experiments. (**p < 0.01; ***p < 0.001; ****p < 0.0001; n.s., not significant).

In addition to YAP/TAZ, we examined the subcellular localization of NF-κB p65, which mediates DNA damage–induced survival signaling through regulation of anti-apoptotic (e.g., Bcl-2, IAPs) and cyclin family genes ^(62)^. NF-κB p65 was analyzed because it transcriptionally links DNA damage responses to cell survival and proliferation independently of Hippo pathway regulation. Immunofluorescence analysis revealed a similar tendency of the NF-κB p65 nuclear localization in both round and elongated nuclei (Fig. 6E). Quantitative nuclear-to-cytoplasmic ratio further showed NF-κB p65 nuclear translocation did not differ significantly between round nuclei and their elongated counterparts (Fig. 6F).

Collectively, these results indicate that altered nuclear shape in hemisphere surface decouples YAP and TAZ nuclear localization while maintaining constitutive NF-κB signaling, suggesting a selective mechanotransduction mechanism under concave geometry that preferentially engages TAZ independently of the other Hippo pathway effector, YAP, and classical inflammatory signaling.

## 3. Discussion

In this study, we explored how geometric confinement affects nuclear shape, mechanotransduction, and epigenetic regulation in SaOS-2 osteosarcoma cells by comparing concave hemispherical PDMS surfaces with flat PDMS controls. Our results show that confining nuclei to a rounder shape has a noticeable impact on nuclear mechanics and subcellular signaling, yet it does not compromise overall cell viability.

First, we engineered concave hemisphere surfaces using GSL, enabling an efficient and highly controllable platform. We achieved higher yield in dimensions between designed and resulting surface. Nuclear morphology was significantly altered within larger diameters particularly in 20 µm diameter. This shape shift led to strong confinement within specific surface diameters, resulting in further phenotypic changes that can be attributed to 3D confinement compared to 2D effects.

We carried out all experiments for a 24-hour period to focus on early nuclear response to mechanical confinement, consistent with previous mechanobiological papers using this timeframe for studying SaOS-2 behaviors ^(63,64)^. Under these conditions, SaOS-2 cells grown on flat PDMS surfaces showed elongated nuclei, which is typical for cells settling normally on a flat surface. In contrast, nuclei confined within concave hemispherical surfaces resulted in rounder shape. According to our computational simulations, mechanical loads generated by cytoskeletal tension were dissipated more uniformly in round nuclei compared to elongated nuclei, supporting earlier findings that nuclear shape affects how stress is distributed within the nucleus ^(65–67)^. Interestingly, even though elongated nuclei accumulated more stress, round nuclei confined in concave surfaces exhibited greater division signals. These findings imply that nuclear rounding under geometric confinement may influence cell division and the success of karyokinesis as reflected by the well-known proliferation marker—Ki67 (*see SI*, Fig. S4A-B), potentially alleviating higher rates of abnormal division _motifs_ ^(68–70).^

Consistent with this, cells within hemisphere surfaces have round nuclei and exhibit stronger Ki67 mean intensity level and more frequent nuclear division, suggesting that mechanical confinement can increase the local concentration of proliferative factors. (*see SI*, Fig. S4C-D). In addition to mean intensity, Ki67 exhibits distinct foci organization associated with heterochromatin organization ^(71)^. In accordance with this notion, the overall organization of Ki67 foci was comparable between the two conditions, although round nuclei tended to display a greater number of smaller foci (*see SI*, Fig. S4E-F). In addition, the average Ki67 fluorescence intensity remained higher in round nuclei than in elongated nuclei, even when the foci organization was grouped by having many or few foci based on their number (*see SI*, Fig. S4G-H). Although the average fluorescence intensity reflects the local concentration of Ki67 signal, the higher mean intensity observed in round nuclei, along with the increased frequency of nuclear division on the patterned substrate, suggests a tendency toward enhanced cell division and changes in heterochromatin organization. In terms of cell survival, overall cell viability assessed by absorbance as a measure of subcellular redox activity, was surprisingly similar across all conditions. This indicates that the changes we observed are probably driven by mechanotransduction, rather than by differences in cell survival ^(72)^. Although the nuclear-to-cytoplasmic ratio of cleaved caspase-3 was slightly higher in round nuclei, the difference was minimal, indicating that mechanical confinement does not appear to reach apoptotic threshold. This suggests that mechanical confinement does not drive cells toward a strong apoptotic fate, consistent with previous reports indicating that acute changes in nuclear morphology induced by external perturbations can modulate growth signals without necessarily activating programmed cell death at certain time points ^(35,73)^. Nonetheless, it is important to note that the influence of nuclear mechanics on the regulation of cell proliferation is inherently multifaceted and complex.

Our results indicate that round nuclei on the hemispherical surfaces, which have lower levels of Lamin A/C, are likely softer and compliant than their elongated counterparts, which show higher Lamin A/C and correspondingly stiffer nuclei. Lamin A/C is known to contribute to nuclear stiffness, in agreement with the findings from our simulations ^(18,19,74)^. Our data revealed that changes in nuclear shape by concavity differentially regulates the level of Lamin A and Lamin C isoforms. Morphological shift was accompanied by a reduction in Lamin A/C intensity, whereas Lamin A levels remained largely unaffected. These findings suggest that Lamin C is more sensitive to nuclear shape and mechanical confinement than Lamin A which seems to be maintaining baseline nuclear integrity. This observation is consistent with studies showing that Lamin C contributes more as a dynamic regulator to changes in nuclear integrity and responds to mechanical perturbations ^(75)^. In the present study, we focused primarily on Lamin A/C due to its well-established mechanosensitivity. Therefore other major perinuclear protein Lamin B1 was not assessed, since as reported previously, lamin A/C respond to nuclear rigidity changes more than B1 type Lamin ^(76,77)^. Concomitantly, nuclei located on hemispherical surfaces exhibited a higher degree of morphological abnormalities, such as irregular nuclear shape or aggregation/fragmentation, compared to those on flat surfaces. Such apparent deformation likely reflect the inherent Lamin A/C characteristic of osteosarcoma cells, rather than curvature-induced alterations ^(78,79)^. Substrate curvature may instead amplify the mechanical susceptibility of nuclei with aberrant lamin organization, making their structural instability more obvious within curved geometries (*see SI*, Fig. S2A, Fig. S2C). Moreover, nuclei with abnormal shapes on hemispherical surfaces exhibited lower mean intensities of Lamin A and Lamin A/ C than those on flat surface (*see SI*, Fig. S2B, Fig. S2D). Furthermore, nuclei on both flat and hemispherical surfaces showed a comparable tendency for lamina wrinkling (data not shown). Round nuclei with lower Lamin A/C levels and more prevalent nuclear fragmentation, accompanied by elevated Ki67 intensity level and a higher frequency of nuclear division may reflect curvature-induced nuclear mechanics akin to those traits inside the tumor. These features might be associated with aggressive-like growth tendency of osteosarcoma tumors as previously reported ^(78–80)^. This point should be considered as a possible explanation rather than a solid conclusion. Nevertheless, these findings contribute to our understanding of the intricate relationship between nuclear morphology and key structural proteins in the nuclear envelope.

One of the main findings in this study is that mechanically confined round nuclei exhibited higher mean H3K9me3 fluorescence than their elongated counterparts, despite no detectable difference in HDAC1 abundance. This suggests that nuclear rounding alone can promote heterochromatin compaction rather than changing global histone deacetylation. Because of the difficulty in accurately detecting discrete H3K9me3 foci due to signal overlap with adjacent regions, we quantified MFI rather than foci. As foci become more condensed, their increased brightness is reflected in higher mean intensity values, providing a reliable measure of H3K9me3 levels. Changes in nuclear morphology are likely to have a selective impact on this chromatin modification, because H3K9me3-enriched heterochromatin is strongly anchored with the lamina-associated domains, thereby amplifying the observed effect ^(81–85)^. These findings align with earlier research demonstrating that transcriptional activity and heterochromatin organization can be influenced by nuclear architecture ^(22,86–88)^. This notion is supported by our findings, which shows that mechanically driven reductions in lamin A/C level weaken chromatin tethering at the nuclear periphery, thereby enabling reorganization of heterochromatin to condense more efficiently in a round nucleus. Recent research further strengthened that chromatin and lamin A/C are dynamic regulators of nuclear stiffness under slight deformations ^(87,89)^. While our 24-hour time point captures a relevant mechanoadaptive state associated with altered H3K9me3 levels and nuclear morphology, longer confinement durations may reveal additional chromatin adaptations or recovery phases not observable at a single time point ^(22,90)^. This understanding supports the idea that the observed elevated H3K9me3 in mechanically confined round nuclei reflects a stable mechanotransductive chromatin adaptation at 24 h without substantial alterations on the global epigenetic landscape. Thus, longer time observations are needed to fully characterize the progression of nuclear shape–lamin–H3K9me3 axis and its phenotypic impacts for future investigation. Overall, this observation highlights the importance adaptive response of nuclear rounding within concave geometry in safeguarding genome integrity to prevent aberrant transcription through chromatin compaction.

We assessed DNA damage by comparing average γH2AX fluorescence levels in cells with altered nuclear morphology. Although γH2AX levels were slightly higher in cells with rounded nuclei, they were still substantially lower than the levels observed following irradiation-induced DNA damage. It implies that mechanical confinement induces only minimal,physiological replication stress rather than pathological DNA damage. In this regard, the detected DNA damage was likely subthreshold and repairable, consistent with apoptosis results implying a non-lethal stress response rather than catastrophic genotoxicity (Fig. 5C-D). As a matter of fact, previous studies have shown that altered nuclear shape can increase DNA tension and replication stress causing DNA damage under mechanical confinement ^(22,91)^. Furthermore, the modest increase in γH2AX and a concurrent elevation in H3K9me3 level may seem contradictory, as H3K9me3-enriched heterochromatin is mostly correlated with genome stability and transcriptional repression. However, the observed pattern in γH2AX and H3K9me3 axis shows an adaptive-protective response rather than a failure of genome maintenance. Increased H3K9me3 has been reported to function as a chromatin-based buffering mechanism that limits the extent of DNA damage ^(92–94)^. From this perspective, heterochromatin does not act as an absolute shield, but instead keeps damage within a tolerable/sublethal range. More specifically, H3K9me3-marked heterochromatin reduces chromatin accessibility and locally tightening the genome, thereby limiting aberrant transcription and further defects. When this heterochromatin-mediated adaptation is sufficient, γH2AX accumulation remains modest. On the other hand, failure of this protective response is likely to develop more pronounced DNA damage signal ^(92,95)^. Together, these observations support a model in which mechanically induced chromatin remodeling—particularly increased H3K9me3—acts as an adaptive and protective mechanism to preserve genome integrity under physical confinement.

Two major effectors in Hippo pathway—YAP and TAZ, were found to respond differently depending on alteration of nuclear morphology within hemisphere surfaces. Consistent with previous studies showing that nuclear flattening favors YAP nuclear localization ^(96,97)^, we observed that YAP was predominantly retained in the cytosol of rounded nuclei. Unlike YAP, TAZ showed a stronger tendency to accumulate in the round nuclei compared to elongated ones. Though these two pillars of Hippo pathway are often redundant, they can be divergent with distinct targets as previously indicated ^(98,99)^. This decoupled pattern suggests a subtle, non-canonical, geometry-dependent regulation of YAP and TAZ. Intriguingly, both YAP and TAZ also displayed a greater tendency for nuclear localization on soft substrates. Classical studies suggested that YAP/TAZ preferentially localizes to the nucleus on stiff substrates, largely through actomyosin-driven mechanisms ^(4)^. However, subsequent work revealed that it is the transmission of forces to the nucleus itself that regulates YAP nuclear entry ^(96)^. More recently, it has been shown that compressing or deforming the nucleus is sufficient to permit YAP nuclear entry, partially independent of the cytoskeleton and substrate stiffness ^(100)^. Together, these findings highlight how substrate stiffness, cytoskeletal forces, and nuclear shape can act independently, explaining why YAP/TAZ can occasionally enter the nucleus even on soft surfaces. Although we did not measure F-actin abundance, the higher nuclear accumulation of YAP in elongated nuclei may reflect earlier findings that YAP activation depends heavily on cytoskeletal tension ^(4,96,97,101)^. Correspondingly, higher abundance of TAZ in round nuclei likely act as a compensatory regulator under these conditions.

Meanwhile, we also investigated NF-κB p65, whose nuclear-to-cytoplasmic ratio remained unchanged between round and elongated nuclei. Mechanical cues have previously been shown to cooperate with inflammatory pathways, enhancing NF-κB nuclear translocation in response to cytoskeletal stress ^(102,103)^. Although osteosarcoma cells constitutively activate NF-κB signaling even in the absence of external stimuli such as lipopolysaccharide (LPS) (Fig. 6F; *see SI*, Fig. S6) ^(104)^, our results indicate that pro-inflammatory signaling in SaOS-2 cells is not affected by nuclear mechanical confinement. In parallel, the modest increase in γH2AX levels observed in round nuclei within hemisphere surface, which also exhibited lower lamin A/C, is consistent with the notion that reduced lamin A/C levels favor curvature-driven damage and γH2AX accumulation, as reported by *Xia et al.*,^(25)^. This pattern is compatible with sublethal replication stress which still permits cell-cycle progression and cell survival (Fig. 5; *see SI*, Fig. S4) ^(105)^. In canonical DNA damage response pathway, elevated γH2AX levels can activate NF-κB transcription factor subunit p65 via the IκB kinase linking to ATM kinase to promote cell survival without engaging p53-dependent cell cycle arrest ^(106–108)^. The p53-null status of SaOS-2 cells provides an important contextual framework for interpreting these findings ^(109)^. Therefore, the slight enrichment of γH2AX in confined nuclei may reflect damage-tolerant state in SaOS-2 cells lacking p53 consistent with an adaptive and distinct growth phenotype^(110)^. This behavior supports damage-tolerant growth often seen in aggressive tumor cells ^(111,112)^. Together, these results suggest that the bone niche shapes how tumor cells reorganize subnuclear processes to fit within the confined architecture of Howship’s lacunae, beyond classical stress or inflammatory signaling ^(113)^.

This study models osteosarcoma microenvironment by mimicking Howship’s lacunae concavities, distinct from many stiffness-driven approaches. Future whole-cell geometric confinement platforms, combined with time-lapse imaging and genome-wide transcriptomics, will enable the formulation of causal conclusions, the identification of YAP/TAZ downstream targets, and the revelation of dynamic, paralog-specific mechanotransductive programs underlying tumor plasticity.

## 4. Conclusions

In summary, concave geometry imposing nucleus confinement induces a nuclear rounding. Such change in nuclear shape reprograms mechanical and epigenetic state with largely preserved viability. This geometry-specific adaptive response in osteosarcoma cells redistributes mechanical stress evenly along the nucleus, reduces lamin A/C levels, and induces chromatin compaction with elevated H3K9me3 levels. These changes result in sublethal DNA damage that may prevent pathological genomic instability while activating mechanosensitive pathways facilitating TAZ nuclear translocation without triggering apoptosis. This study shows how substrate curvature reprograms osteosarcoma nuclear mechanobiology, driving lamina remodeling, chromatin reorganization, growth pattern and differential YAP/TAZ signaling without affecting canonical inflammatory signaling. This axis supports the idea that nuclear geometry can act as an important regulator of subnuclear dynamics, providing a notion for understanding how the microenvironment mimicking trabecular bone or Howship’s lacunar architecture influence subnuclear dynamics in cancers, particularly those with poor prognoses associated with high nuclear shape plasticity.

## 5. Material and Methods

### Substrate Fabrication and PDMS Molding

Hemisphere surfaces were fabricated using grayscale lithography (GSL), a precise, time-efficient and highly controllable method for generating microconcave structures. The comprehensive methodology including surface calibration and characterization has been detailed in the previous paper ^(50)^.

Initially, master molds were prepared by coating glass slides with a layer of photoresist. Standard microscope glass slides were cut into 2 cm × 2 cm squares and cleaned sequentially by sonication in soapy distilled water, acetone, and isopropanol for 2 minutes each. The cleaned slides were then air-dried to remove any remaining residuals and ensure a clean surface for further steps. Following completion of the cleaning protocol, each glass substrate was uniformly coated with a 40 µm layer of ma-P 1275G photoresist (Micro Resist Technology) through spin coating at 450 rpm for 40 seconds. Thereafter, the photoresist-coated substrates were subjected to thermal treatment on a hotplate at 100 °C for 10 minutes to ensure effective cross-linking and to mitigate nitrogen bubble formation within the film.

Before UV exposure, a customized digital mask design protocol for GSL was implemented. Mask image files were generated in the open-source software Blender (blender.org), which, although mainly used for visualization and design applications ^(114)^, it was adapted here for precise construction of 3D microstructures, assignment of height-based color gradients and orthogonal rendering to remove perspective distortion. The hemispherical features were created in Blender and arranged into a honeycomb appearance array. Height-based shading was achieved using a calibration curve that mapped Z-axis values to corresponding color gradients. The resulting orthogonally rendered images were used as calibrated digital masks for mold fabrication.

Photoresist exposure was conducted using a Smart Print UV lithography system (Microlight 3D) equipped with a ×10 objective lens. The substrates were illuminated for five seconds at 20 % UV intensity with a 385 nm light source (approximately 1920 mW/cm²). After exposure, the films were developed in mr-D 526/S developer (Micro Resist Technology, Germany) for three minutes, enabling selective dissolution of carboxylic acid groups and revealing the desired microstructures. No post-exposure bake was applied. The resulting microstructured molds were characterized by Scanning Electron Microscopy (Fei Quanta400, China), confocal microscopy (LSM 800, Zeiss, USA), and profilometer (Bruker, Germany) to assess the fidelity and dimensional accuracy of the fabricated features following the previously established workflow.

The master molds were then proceeded for replication of the hemisphere surfaces through polydimethylsiloxane (PDMS) molding. A mixture of SYLGARD™ 184 Silicone Elastomer Kit (Dow Corning, USA), prepared at a 10:1 ratio of prepolymer to crosslinking agent, was poured over the master molds. The PDMS was cured at 70 °C for 2 hours to ensure complete crosslinking, after which the replicas were carefully demolded to obtain the final PDMS pieces containing hemisphere surfaces. An aqua-silicone sealant (Den Braven, the Netherlands) was used to |attach the clean PDMS piece containing the hemisphere design to either 35 mm or 60 mm tissue culture petri dishes. Samples were cleaned with a two-step wash (70% ethanol and distilled water) before being sterilized with UV light for 15 minutes. Soft PDMS samples were also prepared by mixing silicone elastomer base with a curing agent at a 50:1 ratio (w/w) for YAP/TAZ observations. Before cell seeding, PDMS samples either 10:1 or 50:1 were functionalized with fibronectin from bovine plasma (50 µg/ml, Sigma, USA) at room temperature for 40 minutes.

## Cell Culture and Treatments

The SaOS-2 human osteosarcoma cells (ATCC, USA) were maintained in L-glutamine fortified McCoy’s 5 A Modified Medium (Gibco, USA) supplemented with Penicillin-Streptomycin (1% v/v, Sigma, USA) and fetal bovine serum (15% v/v, Gibco, USA) at 37°C and 5% CO_2_ in a humidified incubator. SaOs-2 cells were cultured with no more than 30 passage numbers. Cells were seeded on fibronectin-functionalized hemisphere PDMS molds and cultured for 24 hours in a humidified incubator. Cells were seeded in either a 35 mm or 60 mm petri dish at a density of 10000 cells per cm^2^ and multiplied by 8.8 or 21.5 depending on the petri dish size.

To evaluate γH2AX induction, control cells were exposed to UV irradiation for 20 or 40 seconds using a 254 nm UV lamp equipped with a container (UVP, USA). Following exposure, cells were returned to standard culture conditions (37 °C, 5% CO₂) to allow foci formation for 90 minutes before fixation and immunostaining, as previously described ^(115)^. For NF-κB activation controls, cells were treated with lipopolysaccharide (LPS; 1 µg/mL; Invitrogen, USA) in complete culture medium for 1, 2, or 6 hours to promote NF-κB nuclear translocation, in accordance with earlier reports ^(116)^. During the treatment period, cells were incubated at 37 °C in a humidified atmosphere containing 5% CO₂ before fixation and immunostaining. Cleaved caspase-3 level was induced by nutrient and serum deprivation ^(117)^. Cells were treated with Hanks’ Balanced Salt Solution (HBSS, Sigma, USA) for 24 hours at 37 °C in a humidified 5% CO₂ incubator, after which immunostaining was performed.

## Cell Viability Assay

Before proceeding with the experiment, PDMS hemispherical replicates and PDMS flat pieces were assembled into 24 wells and handled as indicated in the previous section (*see Substrate Fabrication, PDMS Molding and Surface Functionalization*). After the well plate had been prepared, 30 000 cells in 500 µl were plated in each well onto surfaces coated with fibronectin (50 µg/ml) for each time-course observation (24 and 48 hours). Polystrene wells without PDMS pieces and cell medium without suspension of cells were used as control groups. The cells were then incubated overnight at 37°C and 5% CO_2_ in a humidified incubator. Following aspiration of the culture medium, alamarBlue reagent (10% (v/v), Invitrogen) was prepared in 500 µl fresh medium and introduced with the tested conditions. Samples were incubated for 6 hours at 37°C and 5% CO_2_ and then measured with a plate reader (Multiskan FC, Thermoscientific) at 570 nm and 595 nm. All readouts were calculated using the following formula:

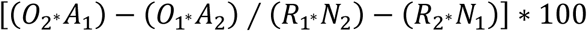

*O1* = molar extinction coefficient of oxidized alamarBlue (blue) at 570 nm
*O2* = molar extinction coefficient of oxidized alamarBlue at 600 nm
*R1* = molar extinction coefficient of reduced alamarBlue (red) at 570 nm
*R2* = molar extinction coefficient of reduced alamarBlue at 600 nm
*A1* = absorbance of tested wells at 570 nm
*A2* = absorbance of tested wells at 600 nm
*N1* = absorbance of negative control well (medium + alamarBlue without cells) at 570 nm
*N2* = absorbance of negative control well (medium + alamarBlue without cells) at 600 nm

## Immunofluorescence

Immunofluorescence staining was performed according to standard procedures by Thermo Fischer Scientific. After 24 hours of incubation, the cells on surface were fixed in 4% paraformaldehyde (PFA) in 1X phosphate-buffered saline (PBS) for 10–15 minutes at 37 °C, followed by PBS washing for three times. Samples were then permeabilized with 0.1% Triton X-100 in 1X PBS for 15 minutes at room temperature to facilitate subcellular labeling and washed three more times with PBS. To minimize non-specific binding, the samples were blocked with 2% bovine serum albumin (BSA) for 1 hour at room temperature, followed by three PBS washes.

Samples were incubated with primary antibodies rabbit anti-Lamin A (2 μg/ml, L1293, Sigma-Aldrich), rabbit anti-Lamin A/C (1:250, ab108595, Abcam), rabbit anti-Histone Deacetylase 1 (HDAC1, 1:500, H3284, Sigma-Aldrich), rabbit anti-Histone H3 trimethyl K9(H3K9me3, 0.5 μg/ml, ab8898, Abcam), rabbit anti-γ-H2AX phospho S139 (1:250, ab81299, Abcam), rabbit anti-Cleaved Caspase-3 Asp175 (1:400, 9661, Cell Signaling Technology), rabbit anti-YAP D81HX (1:100, 14074, Cell Signaling Technology), rabbit anti-TAZ (1:100, ab84927, Abcam), mouse anti-NF-κB p65 (2 μg/ml, ab13594, Abcam) diluted in 0.1% BSA in 1× PBS, at the concentrations recommended by the manufacturer for 3 h at room temperature. Since the manufacturer no longer produces the anti-TAZ and anti-NF-κB p65 antibodies in these catalog numbers, the final concentrations of both antibodies were determined based on the recommended dilutions of currently available antibodies from the same manufacturer (anti-TAZ: ab277791; anti-NF-κB p65: ab16502). After extensive washing with PBS, samples were incubated with appropriate fluorophore-conjugated secondary antibodies depending on staining strategy including Anti-Rabbit IgG-TRITC (1:150, T6778, Sigma-Aldrich), Anti-Rabbit IgG-FITC (1:150, F9887, Sigma-Aldrich), Anti-Mouse IgG-FITC (1:500, F0257, Sigma-Aldrich) diluted in 0.1% BSA in 1× PBS for 1 h at room temperature in the dark. Nuclei and the actin cytoskeleton were counterstained with bisBenzimide Hoechst 33342 (20 μg/mL, B2261, Sigma-Aldrich) and either Alexa Fluor™ 488 (1:75, A12379, Thermo Fisher Scientific) or Alexa Fluor™ 555 (1:75, A34055, Thermo Fisher Scientific) or Alexa Fluor™ 647 (1:300, A30107, Thermo Fisher Scientific), respectively. The staining probes were diluted in 0.1% BSA in 1× PBS as applied concurrently with the secondary antibody staining for 1 h at room temperature keeping the samples protected from light. To assess cell proliferation, fluorescently conjugated rabbit Ki67 (10 μg/ml; 14-5698-82, Thermo Fisher) was diluted in 0.1% BSA in 1× PBS and incubated for 1 hour at room temperature in the dark, allowing proper staining without the need for a primary antibody. Following three additional PBS washes, samples were resuspended in 1× PBS and stored at 4 °C for up to 3–4 days prior to imaging.

## Image Acquisition

Fluorescence imaging was performed using a confocal laser scanning microscope (LSM 800, Zeiss, USA) equipped with a ×63 water/PBS-immersion lens. Pinhole was set to be 1 Airy unit for all channels. Cells were imaged on fibronectin-coated PDMS substrates. Additional controls including starvation medium, LPS treatment, and irradiation were indicated relevant section (*see Cell culture and treatments*). 3D images were acquired with a z-slice scaling of 0.4 µm, covering the full nuclear volume for all conditions. Optical zoom was kept constant within individual experiments; 2× digital zoom was used where higher spatial resolution of nuclear structures was required at a frame size of 512×512 pixels, while no zoom was applied for whole-cell analyses at a frame size of 1024×1024 pixels. Laser lines appropriate for each fluorophore were used according to manufacturer recommendations. Laser power and detector gain were kept identical within each experimental repetition and condition, but were varied between different experimental sets to optimize signal-to-noise ratios for specific antibodies and fluorophores (Table 1). Importantly, all quantitative comparisons were performed between samples imaged using identical acquisition settings based on experimental set-up, and no comparisons were made across images acquired with different gain or laser settings. Images were acquired below saturation levels, as verified using the Zeiss range indicator. Raw images were used for quantitative analysis. Only linear adjustments of brightness and contrast were applied for figure presentation, and these adjustments were applied equally across compared images.

**Table 1.** Confocal microscope acquisition settings of each fluorophore for different experiments

| Experiment Name or Purpose | 405 nm Detector Gain&Laser Wavelength | 488 nm Detector Gain&Laser Wavelength | 555 nm Detector Gain&Laser Wavelength | 640 nm Detector Gain&Laser Wavelength |
| --- | --- | --- | --- | --- |
| Nucleus Shape | 700 V-75 % | - | 750 V-75 % | - |
| Lamin A | 750 V-50 % | 700 V-5 % | - | - |
| Lamin A/C | 850 V-25 % | 700 V-5 % | - | - |
| HDAC1 | 750 V-35 % | 750 V-5 % | - | - |
| H3K9me3 | 750 V-35 % | 700 V-5 % | - | - |
| Ki67 | 850 V-50 % | 800 V-3 % | - | - |
| γH2AX | 800 V-50 % | 850 V-10 % | - | - |
| C-Caspase 3 | 800 V-50 % | 750 V-1.5 % | 800 V-7 % | - |
| YAP | 850 V-50 % | 750 V-1 % | 750 V-10 % | - |
| TAZ | 850 V-35 % | 850 V-5 % | - | 750 V-0.2 % |
| NF-kB | 800 V-50 % | 850 V-10 % | - | 700 V-0.3 % |

Surface topography and elastic modulus of the PDMS surfaces (1:10 and 1:50) were determined by Atomic Force Microscopy (AFM). AFM measurements were performed using a Bruker Multimode IV, with a Nanoscope V controller and an E “vertical” scanner (Bruker), by the Peak Force Quantitative Nanomechanical Mapping (PF-QNM, Bruker) method ^(118)^. PF-QNM is a contact AFM mode, based on the force-volume method. A calibration procedure was first followed which consist of precise measurements of the tip’s radius, spring constant, and resonance frequency. All quantitative measurements were carried out with NCLR cantilever (Nano World) with a spring constant of 48 N/m and a resonance frequency of 190 kHz, a width of 38 μm and a length of 225 μm. Thanks to the Sader method ^(119)^, the actual spring constant was determined and found to be around 27 N/m. Then, the deflection sensitivity (around 32 nm/V) was measured on a sapphire surface. The Tip radius was calibrated against a polystyrene standard provided by Bruker. The measured value of the tip radius was 10 nm. The Poisson’s ratio was assumed to be equal to 0.3. To get relevant results, the cantilever and the tip geometry are taken into account in the PF-QNM measurements. For all experiments, an area of 1 μm × 1 μm (256 × 256 pixels at 0.6 Hz) were taken at three different areas on the sample surface.

## Image analysis

For quantification of nuclear fluorescence intensity, z-stack images of nuclei channels were projected as sum intensity projections using ImageJ (Fiji). Before nuclear segmentation, the projected nuclear images were preprocessed using an edge-preserving median filter (radius = 5.0) to reduce noise and improve thresholding accuracy. Z-stack images of the target protein were projected into a single image using sum-intensity projection prior to mean fluorescence intensity (MFI) measurements. Nuclear regions of interest were defined using a DAPI mask, and the same nuclear masks were applied into the target protein channel for accurate comparison.

3D nuclear shape and volume analyses were performed using Imaris software. Z-stack images were imported into Imaris, and nuclei were segmented using the Surfaces module and 3D surface reconstructions were generated for each nucleus, from which nuclear volume and shape parameters (including sphericity and surface area) were quantified automatically. For 2D analyses, nuclear morphology was quantified in ImageJ using standard shape descriptors, including nuclear area, circularity (calculated from area and perimeter), aspect ratio, and roundness (calculated based on aspect ratio).

Nuclear depth was determined directly from the acquired z-stacks using ZEISS ZEN software by measuring the axial extent of each nucleus from the first to the last z-plane in which the DAPI signal was detected.

Nuclear-to-cytoplasmic (N/C) ratios were quantified from immunofluorescence images using ImageJ/Fiji following a previously described approach ^(120)^. The nuclei were identified using DAPI staining, and the overall cell boundaries were defined based on phalloidin staining of F-actin. Nuclear regions of interest (ROIs) were defined by applying a threshold to the DAPI channel, while whole-cell ROIs were drawn by thresholding the phalloidin channel. The cytoplasmic ROI was defined by subtracting the nuclear ROI from the corresponding whole-cell ROI. Z-stack images of the target protein were projected into a single image using sum-intensity projection prior to MFI measurements. MFI of the target protein was measured separately within nuclear and cytoplasmic ROIs. The N/C ratio was calculated by dividing nuclear MFI by cytoplasmic MFI.

Nuclear foci were quantified using a workflow slightly modified from previously published methods ^(121,122)^. For each nucleus, the target protein channel was projected as a maximum-intensity projection. The resulting images were processed using an unsharp mask (radius = 10, mask weight = 0.7) to enhance foci contrast, followed by Gaussian blur (σ = 0.2, scaled units) to reduce noise. Foci were segmented using Otsu automatic thresholding. Binary images were subsequently processed by filling holes and applying watershed separation to resolve adjacent foci. Individual foci were then quantified using particle analysis within nuclear regions.

## Modelling

To gain further insight into how hemispheric patterns influence mechanical stress on the SaOS-2 nucleus during self-confinement, we employed a mechanical model of an adherent cell developed by *Vassaux et al.* (*2017, 2018*) ^(123,124)^. All simulations were performed using the LMGC90 software package as previously used ^(125)^, written in Python, with additional custom scripts developed to manage the simulation workflow.

The SaOS-2 cell is simulated using a granular tensegrity-based mechanical framework in which the cell and its components including nucleus, cytoskeleton, cytosol, and membrane are represented as nodes and particles interacting through contacts, elastic springs, and compressive cables. This network of reticulated mechanical interactions reproduces the prestressed filamentous architectures of the cytoskeleton and nucleoskeleton embedded within the cytosol and nucleosol. The mechanical state of the SaOS-2 cell model emerges from a balance between contractile forces generated by the actomyosin network and compressive forces arising from microtubules, intermediate filaments, and the cytosol. In the cell model, the nucleus is represented as a dense packing of interacting particles capable of reorganization and behaving as a viscous incompressible fluid, enclosed by a stretchable envelope modeling the nuclear lamina. Nuclear resistance to compressibility arises from the mechanical coupling between the elastic lamina and the nucleoplasm. Nuclear compressibility therefore results from rearrangements of nucleoplasmic particles constrained by lamina stiffness.

To quantitatively reproduce the compliant nuclear state observed in SaOS-2 cells cultured on hemispherical substrates, we adjusted both the stiffness of tensile elastic interactions between lamina nodes and the stiffness of compressive elastic interactions between nucleoplasmic particles. The resulting mechanical properties of the modeled nucleus were characterized by simulating tip-less AFM compression experiments performed in vitro on isolated nuclei, following the approach described previously ^(126)^. The tangent elastic modulus was determined as the slope of the stress–strain curve at 20% compression. Virtual AFM compression simulations yielded elastic moduli of 0.22 kPa for isolated nuclei and 0.35 kPa for nuclei within intact SaOS-2 cell models.

Cell adhesion to hemispherical topographies is simulated in two steps. First, the cell is generated in an elliptical configuration and spread onto the substrate. Second, it is allowed to relax for a sufficiently long time to reach mechanical equilibrium. The model configuration used in this study is identical to that previously validated and calibrated against experimental data by *Wakhloo et al.* (*2020*) ^(45)^. The model does not include active regulatory dynamics of the cytoskeleton or nucleoskeleton. Instead, it isolates the contributions of adhesion conditions, cytoskeletal organization, and cytoskeleton–nucleus mechanical coupling to nuclear deformation.

## Statistical Analysis

Unless stated otherwise, data are presented as median with interquartile range. Statistical analyses were performed using nonparametric tests—either Kruskal–Wallis or Mann–Whitney—depending on sample size, unless specified otherwise. Statistical significance was defined as a p-value below 0.05, with higher levels of significance indicated by **: p ≤ 0.01, ***: p ≤ 0.001, and ****: p ≤ 0.0001. All statistical analyses and graphical representations were generated using GraphPad Prism. Each experiment was performed independently with three biological replicates. Graphs were redrawn by Affinity Designer for layout refinement and structural consistency. A single illustration was created by Biorender.

## Abbreviations

AFM: atomic force microscopy
BSA: bovine serum albumin
GSL: grayscale lithography
HDAC1: histone deacetylase 1
H3K9me3: histone H3 trimethyl K9
LPS: lipopolysaccharide
MFI: mean fluorescence intensity
PFA: paraformaldehyde
TAZ: PDZ-binding motif
PBS: phosphate buffered saline
PDMS: polydimethylsiloxane
ROI: region of interest
SI: supplementary information
3D: three dimensional
2D: two dimensional
YAP: Yes-associated protein

## Acknowledgments

The authors gratefully acknowledge all members of the Biointerface Group at IS2M for their valuable feedback and constructive comments with special thanks to Dr. Carole Arnold, Dr. Isabelle Brigaud and Dr. Nathalie Couturier for their particular contributions. Moreover, we cordially thank Prof. Nan Ma and her team and Prof. Katia Carneiro for insightful scientific discussions.

## Authors contributions

**IT:** Investigation; Methodology; Formal analysis; Validation; Data curation; Visualization; Writing – original draft; Writing – review & editing.

**FFB:** Methodology; Investigation; Validation; Formal analysis; Writing – review & editing.

**JLM:** Formal analysis; Software; Investigation; Funding acquisition; Writing – review & editing.

**PK:** Investigation; Methodology; Validation; Writing – review & editing.

**TP:** Methodology; Supervision; Investigation; Writing – review & editing.

**MB:** Methodology; Investigation; Validation; Formal analysis; Writing – review & editing.

**VL:** Funding acquisition; Writing – review & editing.

**KA:** Conceptualization; Supervision; Funding acquisition; Writing – review & editing. **LP:** Conceptualization; Methodology; Supervision; Validation; Funding acquisition; Writing – review & editing.

## Availability of data and materials

The authors confirm that all relevant data are provided in the main manuscript and the supplementary materials. Additional data produced in this study are available from the corresponding authors, I.T. and L.P., upon reasonable request. Research materials associated with this study can also be obtained from L.P. upon request.

---

**Supplementary Figure S1.** 3D and 2D nuclear morphometric parameters on hemispherical surfaces of varying lengths. Quantification of 3D **(A-B)** and 2D **(C-E)** nuclear characteristics of cells on hemispherical surfaces of varying lengths. **(A)** Volume, *(B-C)* Area, **(D)** Perimeter and **(E)** Aspect ratio. Experiments were performed only once. (*p < 0.05; **p < 0.01; ***p < 0.001; ****p < 0.0001; n.s., not significant).

**Supplementary Figure S2.** Nuclear aggregation within hemisphere surface is associated with reduced Lamin A and Lamin A/C levels. Representative DIC, Hoechst (nuclei), Lamin A **(A)**, Lamin A/C **(C)**, and merged images of SaOS-2 cells exhibiting nuclear aggregation/fragmentation. Scale bar, 5 µm. **(B, D)** Quantification of MFI for Lamin A **(B)** and Lamin A/C **(D)** in elongated, round, and aggregated nuclei. Aggregated nuclei show significantly reduced Lamin A and Lamin A/C levels compared to elongated nuclei, while round and aggregated nuclei exhibit comparable Lamin A/C levels. “n” represents the number of nuclei analyzed from one experiment. (***p < 0.001; ****p < 0.0001; n.s., not significant.)

**Supplementary Figure S3.** Quantification of γH2AX MFI in irradiated control samples. “n” represents the number of nuclei analyzed from only one experiment. (n.s., not significant)

Supplementary Figure S4. Hemispherical substrates promote nuclear division and elevated Ki67 levels independently of foci organization.

**(A)** Representative immunofluorescence images showing nuclear division events with DIC, Hoechst (nuclei), and Ki67 (red) in three independent ROIs in SaOS-2 cells cultured on concave hemispherical PDMS substrates (10 µm depth, 20 µm diameter). Dividing nuclei were more frequently observed on hemispherical surfaces. Arrows are indicating extended Ki67 foci between dividing nuclei. Scale bar, 5 µm. **(B)** Quantification of Ki67 mean fluorescence intensity (MFI) in elongated (flat), round (hemisphere-confined), and dividing nuclei, showing elevated levels in round nuclei and lowest in dividing nuclei. “n” represents the number of nuclei analyzed from only one experiment. **(C)** Representative immunofluorescence images of Ki67 in SaOS-2 cells on flat and concave hemispherical PDMS substrates. Ki67 (red) and Hoechst (light blue) for nucleus; corresponding DIC images are shown. Scale bar, 5 µm. **(D)** Quantification of Ki67 MFI revealed a significant increase in round and mechanically confined nuclei compared with elongated nuclei on flat PDMS **(E)** Representative images showing Ki67 foci organization in elongated and round nuclei. Arrows show smaller foci. Scale bar, 5 µm. **(F)** Quantification of Ki67 foci organization calculated as (foci number/nucleus) × (mean foci area/nucleus). No significant difference was detected between elongated and round nuclei. **(G, H)** Ki67 MFI in nuclei with small/numerous **(G)** vs. large/few **(H)** foci; round nuclei consistently show higher MFI than elongated nuclei. **(D-H)** “n” represents the number of nuclei analyzed from three independent experiments. **(B-H)** (*p < 0.05; **p < 0.01; ***p < 0.001; ****p < 0.0001; n.s., not significant).

**Supplementary Figure S5. Mechanical characterization of substrates by AFM** Mean elasticity values (±SD) for polystyrene, 10:1 PDMS, and 50:1 PDMS substrates, reported in MPa or GPa. Representative AFM topography images are shown alongside, with color scale bars indicating surface roughness (darker/lighter brown = lower/higher features, respectively).

Supplementary Figure S6. Quantification of NF-κB p65 MFI nuclear-to-cytosol (N/C) ratio in time-dependent LPS treated SaOS-2 cells Cells were treated with 1 μg/ml LPS at 1, 2, and 6 hours. “n” represents the number of nuclei analyzed from one experiment. (*p < 0.05; n.s., not significant.)

## References

1. Bakhshandeh B, Ranjbar N, Abbasi A, Amiri E, Abedi A, Mehrabi MR, et al. Recent progress in the manipulation of biochemical and biophysical cues for engineering functional tissues. Bioeng Transl Med. 2023 Mar;8(2):e10383. doi:10.1002/btm2.10383 PubMed PMID: 36925674; PubMed Central PMCID: PMC10013802.

2. Lee J, Jeon O, Kong M, Abdeen AA, Shin JY, Lee HN, et al. Combinatorial screening of biochemical and physical signals for phenotypic regulation of stem cell-based cartilage tissue engineering. Sci Adv. 2020 May;6(21):eaaz5913. doi:10.1126/sciadv.aaz5913 PubMed PMID: 32494742; PubMed Central PMCID: PMC7244269.

3. Li J, Liu Y, Zhang Y, Yao B, Enhejirigala null, Li Z, et al. Biophysical and Biochemical Cues of Biomaterials Guide Mesenchymal Stem Cell Behaviors. Front Cell Dev Biol. 2021;9:640388. doi:10.3389/fcell.2021.640388 PubMed PMID: 33842464; PubMed Central PMCID: PMC8027358.

4. Dupont S, Morsut L, Aragona M, Enzo E, Giulitti S, Cordenonsi M, et al. Role of YAP/TAZ in mechanotransduction. Nature. 2011 Jun 8;474(7350):179–83. doi:10.1038/nature10137 PubMed PMID: 21654799.

5. Iskratsch T, Wolfenson H, Sheetz MP. Appreciating force and shape—the rise of mechanotransduction in cell biology. Nat Rev Mol Cell Biol. 2014 Dec;15(12):825–33. doi:10.1038/nrm3903 PubMed PMID: 25355507; PubMed Central PMCID: PMC9339222.

6. Crowder SW, Leonardo V, Whittaker T, Papathanasiou P, Stevens MM. Material Cues as Potent Regulators of Epigenetics and Stem Cell Function. Cell Stem Cell. 2016 Jan 7;18(1):39–52. doi:10.1016/j.stem.2015.12.012 PubMed PMID: 26748755; PubMed Central PMCID: PMC5409508.

7. Alisafaei F, Jokhun DS, Shivashankar GV, Shenoy VB. Regulation of nuclear architecture, mechanics, and nucleocytoplasmic shuttling of epigenetic factors by cell geometric constraints. Proc Natl Acad Sci U S A. 2019 Jul 2;116(27):13200–9. doi:10.1073/pnas.1902035116 PubMed PMID: 31209017; PubMed Central PMCID: PMC6613080.

8. Arnold C, Tahmaz I, Chapon ML, Maayouf H, Luchnikov V, Milan JL, et al. Bending the rules: curvature’s impact on cell biology. BMC Biol. 2025 Oct 6;23(1):296. doi:10.1186/s12915-025-02416-3 PubMed PMID: 41053697; PubMed Central PMCID: PMC12502368.

9. Luciano M, Tomba C, Roux A, Gabriele S. How multiscale curvature couples forces to cellular functions. Nat Rev Phys. 2024 Mar 8;6(4):246–68. doi:10.1038/s42254-024-00700-9

10. He X, Jiang Y. Substrate curvature regulates cell migration. Phys Biol. 2017 May 23;14(3):035006. doi:10.1088/1478-3975/aa6f8e PubMed PMID: 28535145; PubMed Central PMCID: PMC5572487.

11. Pieuchot L, Marteau J, Guignandon A, Dos Santos T, Brigaud I, Chauvy PF, et al. Curvotaxis directs cell migration through cell-scale curvature landscapes. Nat Commun. 2018 Sep 28;9(1):3995. doi:10.1038/s41467-018-06494-6 PubMed PMID: 30266986; PubMed Central PMCID: PMC6162274.

12. Werner M, Blanquer SBG, Haimi SP, Korus G, Dunlop JWC, Duda GN, et al. Surface Curvature Differentially Regulates Stem Cell Migration and Differentiation via Altered Attachment Morphology and Nuclear Deformation. Adv Sci (Weinh). 2017 Feb;4(2):1600347. doi:10.1002/advs.201600347 PubMed PMID: 28251054; PubMed Central PMCID: PMC5323878.

13. Sun B, Xie K, Chen TH, Lam RHW. Preferred cell alignment along concave microgrooves. RSC Adv. 2017;7(11):6788–94. doi:10.1039/C6RA26545F

14. Werner M, Petersen A, Kurniawan NA, Bouten CVC. Cell-Perceived Substrate Curvature Dynamically Coordinates the Direction, Speed, and Persistence of Stromal Cell Migration. Adv Biosyst. 2019 Oct;3(10):e1900080. doi:10.1002/adbi.201900080 PubMed PMID: 32648723.

15. Vassaux M, Pieuchot L, Anselme K, Bigerelle M, Milan JL. A Biophysical Model for Curvature-Guided Cell Migration. Biophys J. 2019 Sep 17;117(6):1136–44. doi:10.1016/j.bpj.2019.07.022 PubMed PMID: 31400917; PubMed Central PMCID: PMC6818171.

16. Leclech C, Cardillo G, Roellinger B, Zhang X, Frederick J, Mamchaoui K, et al. Micro-Scale Topography Triggers Dynamic 3D Nuclear Deformations. Adv Sci (Weinh). 2025 Mar;12(11):e2410052. doi:10.1002/advs.202410052 PubMed PMID: 39873289; PubMed Central PMCID: PMC11923911.

17. Luciano M, Xue SL, De Vos WH, Redondo-Morata L, Surin M, Lafont F, et al. Cell monolayers sense curvature by exploiting active mechanics and nuclear mechanoadaptation. Nat Phys. 2021 Dec;17(12):1382–90. doi:10.1038/s41567-021-01374-1

18. Srivastava LK, Ju Z, Ghagre A, Ehrlicher AJ. Spatial distribution of lamin A/C determines nuclear stiffness and stress-mediated deformation. J Cell Sci. 2021 May 15;134(10):jcs248559. doi:10.1242/jcs.248559 PubMed PMID: 34028539; PubMed Central PMCID: PMC8186481.

19. Buxboim A, Swift J, Irianto J, Spinler KR, Dingal PCDP, Athirasala A, et al. Matrix elasticity regulates lamin-A,C phosphorylation and turnover with feedback to actomyosin. Curr Biol. 2014 Aug 18;24(16):1909–17. doi:10.1016/j.cub.2014.07.001 PubMed PMID: 25127216; PubMed Central PMCID: PMC4373646.

20. Donnaloja F, Carnevali F, Jacchetti E, Raimondi MT. Lamin A/C Mechanotransduction in Laminopathies. Cells. 2020 May 24;9(5):1306. doi:10.3390/cells9051306 PubMed PMID: 32456328; PubMed Central PMCID: PMC7291067.

21. Lele TP, Dickinson RB, Gundersen GG. Mechanical principles of nuclear shaping and positioning. J Cell Biol. 2018 Oct 1;217(10):3330–42. doi:10.1083/jcb.201804052 PubMed PMID: 30194270; PubMed Central PMCID: PMC6168261.

22. Nava MM, Miroshnikova YA, Biggs LC, Whitefield DB, Metge F, Boucas J, et al. Heterochromatin-Driven Nuclear Softening Protects the Genome against Mechanical Stress-Induced Damage. Cell. 2020 May 14;181(4):800–817.e22. doi:10.1016/j.cell.2020.03.052 PubMed PMID: 32302590; PubMed Central PMCID: PMC7237863.

23. Liu W, Padhi A, Zhang X, Narendran J, Anastasio MA, Nain AS, et al. Dynamic Heterochromatin States in Anisotropic Nuclei of Cells on Aligned Nanofibers. ACS Nano. 2022 Jul 26;16(7):10754–67. doi:10.1021/acsnano.2c02660 PubMed PMID: 35803582; PubMed Central PMCID: PMC9332347.

24. Bunner S, Huang K, Shah A, Figueroa S, Lang N, Chu C, et al. Changes in nuclear and actin mechanics from G1 to G2 affect nuclear integrity. Journal of Cell Science. 2026 Jan 19;jcs.264118. doi:10.1242/jcs.264118

25. Xia Y, Ivanovska IL, Zhu K, Smith L, Irianto J, Pfeifer CR, et al. Nuclear rupture at sites of high curvature compromises retention of DNA repair factors. J Cell Biol. 2018 Nov 5;217(11):3796–808. doi:10.1083/jcb.201711161 PubMed PMID: 30171044; PubMed Central PMCID: PMC6219729.

26. Cho S, Vashisth M, Abbas A, Majkut S, Vogel K, Xia Y, et al. Mechanosensing by the Lamina Protects against Nuclear Rupture, DNA Damage, and Cell-Cycle Arrest. Dev Cell. 2019 Jun 17;49(6):920–935.e5. doi:10.1016/j.devcel.2019.04.020 PubMed PMID: 31105008; PubMed Central PMCID: PMC6581604.

27. Cherdyntseva V, Paulson J, González-Acosta D, Ubieto-Capella P, Rodrigues M, Aouami M, et al. Nucleoplasmic Lamin A/C controls replication fork restart upon stress by modulating local H3K9me3 and ADP-ribosylation levels. Nat Commun. 2025 Nov 29;16(1):11239. doi:10.1038/s41467-025-66098-9

28. Callens SJP, Fan D, van Hengel IAJ, Minneboo M, Díaz-Payno PJ, Stevens MM, et al. Emergent collective organization of bone cells in complex curvature fields. Nat Commun. 2023 Mar 3;14(1):855. doi:10.1038/s41467-023-36436-w PubMed PMID: 36869036; PubMed Central PMCID: PMC9984480.

29. Frey K, Brunner M, Curio C, Kemkemer R. Curvature Perception of Mesenchymal Cells on Mesoscale Topographies. Adv Healthc Mater. 2025 Jan;14(3):e2402865. doi:10.1002/adhm.202402865 PubMed PMID: 39659136; PubMed Central PMCID: PMC11773129.

30. van der Putten C, van den Broek D, Kurniawan NA. Myofibroblast transdifferentiation of keratocytes results in slower migration and lower sensitivity to mesoscale curvatures. Front Cell Dev Biol. 2022;10:930373. doi:10.3389/fcell.2022.930373 PubMed PMID: 35938166; PubMed Central PMCID: PMC9355510.

31. Werner M, Kurniawan NA, Korus G, Bouten CVC, Petersen A. Mesoscale substrate curvature overrules nanoscale contact guidance to direct bone marrow stromal cell migration. J R Soc Interface. 2018 Aug;15(145):20180162. doi:10.1098/rsif.2018.0162

32. Spieker CJ, Závodszky G, Mouriaux C, van der Kolk M, Gachet C, Mangin PH, et al. The Effects of Micro-vessel Curvature Induced Elongational Flows on Platelet Adhesion. Ann Biomed Eng. 2021 Dec;49(12):3609–20. doi:10.1007/s10439-021-02870-4 PubMed PMID: 34668098; PubMed Central PMCID: PMC8671278.

33. Bagchi P. Mesoscale simulation of blood flow in small vessels. Biophys J. 2007 Mar 15;92(6):1858–77. doi:10.1529/biophysj.106.095042 PubMed PMID: 17208982; PubMed Central PMCID: PMC1861774.

34. Mandrycky C, Hadland B, Zheng Y. 3D curvature-instructed endothelial flow response and tissue vascularization. Sci Adv. 2020 Sep;6(38):eabb3629. doi:10.1126/sciadv.abb3629 PubMed PMID: 32938662; PubMed Central PMCID: PMC7494348.

35. Xu X, Wang W, Liu Y, Bäckemo J, Heuchel M, Wang W, et al. Substrates mimicking the blastocyst geometry revert pluripotent stem cell to naivety. Nat Mater. 2024 Dec;23(12):1748–58. doi:10.1038/s41563-024-01971-4 PubMed PMID: 39134648; PubMed Central PMCID: PMC11599042.

36. Yang Y, Xu T, Bei HP, Zhang L, Tang CY, Zhang M, et al. Gaussian curvature-driven direction of cell fate toward osteogenesis with triply periodic minimal surface scaffolds. Proc Natl Acad Sci U S A. 2022 Oct 11;119(41):e2206684119. doi:10.1073/pnas.2206684119 PubMed PMID: 36191194; PubMed Central PMCID: PMC9564829.

37. Florencio-Silva R, Sasso GR da S, Sasso-Cerri E, Simões MJ, Cerri PS. Biology of Bone Tissue: Structure, Function, and Factors That Influence Bone Cells. Biomed Res Int. 2015;2015:421746. doi:10.1155/2015/421746 PubMed PMID: 26247020; PubMed Central PMCID: PMC4515490.

38. Schamberger B, Ehrig S, Dechat T, Spitzer S, Bidan CM, Fratzl P, et al. Twisted-plywood-like tissue formation in vitro. Does curvature do the twist? PNAS Nexus. 2024 Apr;3(4):pgae121. doi:10.1093/pnasnexus/pgae121 PubMed PMID: 38590971; PubMed Central PMCID: PMC10999733.

39. Alias MA, Buenzli PR. Modeling the Effect of Curvature on the Collective Behavior of Cells Growing New Tissue. Biophys J. 2017 Jan 10;112(1):193–204. doi:10.1016/j.bpj.2016.11.3203 PubMed PMID: 28076811; PubMed Central PMCID: PMC5232895.

40. Huang G, Hou T, Song D, Meng T. The regulatory networks and mechanisms of bone microenvironment in tumorigenesis and metastasis. J Bone Oncol. 2025 Dec;55:100729. doi:10.1016/j.jbo.2025.100729 PubMed PMID: 41399767; PubMed Central PMCID: PMC12702220.

41. Shoaib Z, Fan TM, Irudayaraj JMK. Osteosarcoma mechanobiology and therapeutic targets. Br J Pharmacol. 2022 Jan;179(2):201–17. doi:10.1111/bph.15713 PubMed PMID: 34679192; PubMed Central PMCID: PMC9305477.

42. Vanderoost J, van Lenthe GH. From histology to micro-CT: Measuring and modeling resorption cavities and their relation to bone competence. World J Radiol. 2014 Sep 28;6(9):643–56. doi:10.4329/wjr.v6.i9.643 PubMed PMID: 25276308; PubMed Central PMCID: PMC4176782.

43. Xiao P, Schilling C, Wang X. Characterization of Trabecular Bone Microarchitecture and Mechanical Properties Using Bone Surface Curvature Distributions. J Funct Biomater. 2024 Aug 22;15(8):239. doi:10.3390/jfb15080239 PubMed PMID: 39194677; PubMed Central PMCID: PMC11355924.

44. Jinnai H, Watashiba H, Kajihara T, Nishikawa Y, Takahashi M, Ito M. Surface curvatures of trabecular bone microarchitecture. Bone. 2002 Jan;30(1):191–4. doi:10.1016/s8756-3282(01)00672-x PubMed PMID: 11792584.

45. Tusamda Wakhloo N, Anders S, Badique F, Eichhorn M, Brigaud I, Petithory T, et al. Actomyosin, vimentin and LINC complex pull on osteosarcoma nuclei to deform on micropillar topography. Biomaterials. 2020 Mar;234:119746. doi:10.1016/j.biomaterials.2019.119746 PubMed PMID: 31945617.

46. Badique F, Stamov DR, Davidson PM, Veuillet M, Reiter G, Freund JN, et al. Directing nuclear deformation on micropillared surfaces by substrate geometry and cytoskeleton organization. Biomaterials. 2013 Apr;34(12):2991–3001. doi:10.1016/j.biomaterials.2013.01.018 PubMed PMID: 23357373.

47. Singh I, Lele TP. Nuclear Morphological Abnormalities in Cancer: A Search for Unifying Mechanisms. Results Probl Cell Differ. 2022;70:443–67. doi:10.1007/978-3-031-06573-6_16 PubMed PMID: 36348118; PubMed Central PMCID: PMC9722227.

48. Kawaguchi K, Kohashi K, Iwasaki T, Yamamoto T, Ishihara S, Toda Y, et al. Prognostic value of nuclear morphometry in myxoid liposarcoma. Cancer Sci. 2023 May;114(5):2178–88. doi:10.1111/cas.15729 PubMed PMID: 36661410; PubMed Central PMCID: PMC10154898.

49. Lu C, Romo-Bucheli D, Wang X, Janowczyk A, Ganesan S, Gilmore H, et al. Nuclear shape and orientation features from H&E images predict survival in early-stage estrogen receptor-positive breast cancers. Lab Invest. 2018 Nov;98(11):1438–48. doi:10.1038/s41374-018-0095-7 PubMed PMID: 29959421; PubMed Central PMCID: PMC6214731.

50. Borghi FF, Bendimerad M, Chapon ML, Petithory T, Vonna L, Pieuchot L. Rapid prototyping of 3D microstructures: A simplified grayscale lithography encoding method using blender. Micro and Nano Engineering. 2025 Mar;26:100294. doi:10.1016/j.mne.2024.100294

51. Nagayama K, Kodama F, Wataya N, Sato A, Matsumoto T. Changes in the intra- and extra-mechanical environment of the nucleus in Saos-2 osteoblastic cells during bone differentiation process: Nuclear shrinkage and stiffening in cell differentiation. J Mech Behav Biomed Mater. 2023 Feb;138:105630. doi:10.1016/j.jmbbm.2022.105630 PubMed PMID: 36565693.

52. Lammerding J. Mechanics of the nucleus. Compr Physiol. 2011 Apr;1(2):783–807. doi:10.1002/cphy.c100038 PubMed PMID: 23737203; PubMed Central PMCID: PMC4600468.

53. Goff MG, Slyfield CR, Kummari SR, Tkachenko EV, Fischer SE, Yi YH, et al. Three-dimensional characterization of resorption cavity size and location in human vertebral trabecular bone. Bone. 2012 Jul;51(1):28–37. doi:10.1016/j.bone.2012.03.028 PubMed PMID: 22507299; PubMed Central PMCID: PMC3371169.

54. Jones SJ, Boyde A. Histomorphometry of Howship’s lacunae formed in vivo and in vitro: depths and volumes measured by scanning electron and confocal microscopy. Bone. 1993;14(3):455–60. doi:10.1016/8756-3282(93)90179-e PubMed PMID: 8363892.

55. Grewal SIS, Jia S. Heterochromatin revisited. Nat Rev Genet. 2007 Jan;8(1):35–46. doi:10.1038/nrg2008 PubMed PMID: 17173056.

56. Delcuve GP, Khan DH, Davie JR. Roles of histone deacetylases in epigenetic regulation: emerging paradigms from studies with inhibitors. Clin Epigenetics. 2012 Mar 12;4(1):5. doi:10.1186/1868-7083-4-5 PubMed PMID: 22414492; PubMed Central PMCID: PMC3320549.

57. Chen J. The Cell-Cycle Arrest and Apoptotic Functions of p53 in Tumor Initiation and Progression. Cold Spring Harb Perspect Med. 2016 Mar 1;6(3):a026104. doi:10.1101/cshperspect.a026104 PubMed PMID: 26931810; PubMed Central PMCID: PMC4772082.

58. Silva FFVE, Padín-Iruegas ME, Caponio VCA, Lorenzo-Pouso AI, Saavedra-Nieves P, Chamorro-Petronacci CM, et al. Caspase 3 and Cleaved Caspase 3 Expression in Tumorogenesis and Its Correlations with Prognosis in Head and Neck Cancer: A Systematic Review and Meta-Analysis. Int J Mol Sci. 2022 Oct 8;23(19):11937. doi:10.3390/ijms231911937 PubMed PMID: 36233242; PubMed Central PMCID: PMC9569947.

59. Braun F, Bertin-Ciftci J, Gallouet AS, Millour J, Juin P. Serum-nutrient starvation induces cell death mediated by Bax and Puma that is counteracted by p21 and unmasked by Bcl-x(L) inhibition. PLoS One. 2011;6(8):e23577. doi:10.1371/journal.pone.0023577 PubMed PMID: 21887277; PubMed Central PMCID: PMC3160893.

60. Hansen CG, Moroishi T, Guan KL. YAP and TAZ: a nexus for Hippo signaling and beyond. Trends Cell Biol. 2015 Sep;25(9):499–513. doi:10.1016/j.tcb.2015.05.002 PubMed PMID: 26045258; PubMed Central PMCID: PMC4554827.

61. Kolahi KS, Donjacour A, Liu X, Lin W, Simbulan RK, Bloise E, et al. Effect of substrate stiffness on early mouse embryo development. PLoS One. 2012;7(7):e41717. doi:10.1371/journal.pone.0041717 PubMed PMID: 22860009; PubMed Central PMCID: PMC3409240.

62. Wang W, Mani AM, Wu ZH. DNA damage-induced nuclear factor-kappa B activation and its roles in cancer progression. J Cancer Metastasis Treat. 2017;3:45–59. doi:10.20517/2394-4722.2017.03 PubMed PMID: 28626800; PubMed Central PMCID: PMC5472228.

63. Beijer NRM, Nauryzgaliyeva ZM, Arteaga EM, Pieuchot L, Anselme K, van de Peppel J, et al. Dynamic adaptation of mesenchymal stem cell physiology upon exposure to surface micropatterns. Sci Rep. 2019 Jun 24;9(1):9099. doi:10.1038/s41598-019-45284-y PubMed PMID: 31235713; PubMed Central PMCID: PMC6591423.

64. Fanelli G, Alloisio G, Lelli V, Marini S, Rinalducci S, Gioia M. Mechano-induced cell metabolism disrupts the oxidative stress homeostasis of SAOS-2 osteosarcoma cells. Front Mol Biosci. 2023;10:1297826. doi:10.3389/fmolb.2023.1297826 PubMed PMID: 38726050; PubMed Central PMCID: PMC11079223.

65. Zhang J, Alisafaei F, Nikolić M, Nou XA, Kim H, Shenoy VB, et al. Nuclear Mechanics within Intact Cells Is Regulated by Cytoskeletal Network and Internal Nanostructures. Small. 2020 May;16(18):e1907688. doi:10.1002/smll.201907688 PubMed PMID: 32243075; PubMed Central PMCID: PMC7799396.

66. Vishavkarma R, Raghavan S, Kuyyamudi C, Majumder A, Dhawan J, Pullarkat PA. Role of actin filaments in correlating nuclear shape and cell spreading. PLoS One. 2014;9(9):e107895. doi:10.1371/journal.pone.0107895 PubMed PMID: 25251154; PubMed Central PMCID: PMC4177564.

67. Lee H, Adams WJ, Alford PW, McCain ML, Feinberg AW, Sheehy SP, et al. Cytoskeletal prestress regulates nuclear shape and stiffness in cardiac myocytes. Exp Biol Med (Maywood). 2015 Nov;240(11):1543–54. doi:10.1177/1535370215583799 PubMed PMID: 25908635; PubMed Central PMCID: PMC4778402.

68. Miller I, Min M, Yang C, Tian C, Gookin S, Carter D, et al. Ki67 is a Graded Rather than a Binary Marker of Proliferation versus Quiescence. Cell Rep. 2018 Jul 31;24(5):1105–1112.e5. doi:10.1016/j.celrep.2018.06.110 PubMed PMID: 30067968; PubMed Central PMCID: PMC6108547.

69. Mouelhi M, Saffon A, Roinard M, Delanoë-Ayari H, Monnier S, Rivière C. Mitosis sets nuclear homeostasis of cancer cells under confinement [Internet]. 2024 [cited 2026 Feb 23]. Available from: https://elifesciences.org/reviewed-preprints/94975v1 doi:10.7554/eLife.94975.1

70. Moriarty RA, Stroka KM. Physical confinement alters sarcoma cell cycle progression and division. Cell Cycle. 2018;17(19–20):2360–73. doi:10.1080/15384101.2018.1533776 PubMed PMID: 30304981; PubMed Central PMCID: PMC6237433.

71. Sobecki M, Mrouj K, Camasses A, Parisis N, Nicolas E, Llères D, et al. The cell proliferation antigen Ki-67 organises heterochromatin. Elife. 2016 Mar 7;5:e13722. doi:10.7554/eLife.13722 PubMed PMID: 26949251; PubMed Central PMCID: PMC4841783.

72. Mohri Z, Del Rio Hernandez A, Krams R. The emerging role of YAP/TAZ in mechanotransduction. J Thorac Dis. 2017 May;9(5):E507–9. doi:10.21037/jtd.2017.03.179 PubMed PMID: 28616323; PubMed Central PMCID: PMC5465147.

73. Roy B, Venkatachalapathy S, Ratna P, Wang Y, Jokhun DS, Nagarajan M, et al. Laterally confined growth of cells induces nuclear reprogramming in the absence of exogenous biochemical factors. Proc Natl Acad Sci U S A. 2018 May 22;115(21):E4741–50. doi:10.1073/pnas.1714770115 PubMed PMID: 29735717; PubMed Central PMCID: PMC6003522.

74. Khan ZS, Santos JM, Hussain F. Aggressive prostate cancer cell nuclei have reduced stiffness. Biomicrofluidics. 2018 Jan;12(1):014102. doi:10.1063/1.5019728 PubMed PMID: 29333204; PubMed Central PMCID: PMC5750055.

75. González-Cruz RD, Sadick JS, Fonseca VC, Darling EM. Nuclear Lamin Protein C Is Linked to Lineage-Specific, Whole-Cell Mechanical Properties. Cell Mol Bioeng. 2018 Apr;11(2):131–42. doi:10.1007/s12195-018-0518-y PubMed PMID: 29755599; PubMed Central PMCID: PMC5943047.

76. Dechat T, Adam SA, Taimen P, Shimi T, Goldman RD. Nuclear Lamins. Cold Spring Harbor Perspectives in Biology. 2010 Nov 1;2(11):a000547–a000547. doi:10.1101/cshperspect.a000547

77. Lammerding J, Fong LG, Ji JY, Reue K, Stewart CL, Young SG, et al. Lamins A and C but not lamin B1 regulate nuclear mechanics. J Biol Chem. 2006 Sep 1;281(35):25768–80. doi:10.1074/jbc.M513511200 PubMed PMID: 16825190.

78. Urciuoli E, Petrini S, D’Oria V, Leopizzi M, Rocca CD, Peruzzi B. Nuclear Lamins and Emerin Are Differentially Expressed in Osteosarcoma Cells and Scale with Tumor Aggressiveness. Cancers (Basel). 2020 Feb 13;12(2):443. doi:10.3390/cancers12020443 PubMed PMID: 32069980; PubMed Central PMCID: PMC7073215.

79. Evangelisti C, Paganelli F, Giuntini G, Mattioli E, Cappellini A, Ramazzotti G, et al. Lamin A and Prelamin A Counteract Migration of Osteosarcoma Cells. Cells. 2020 Mar 22;9(3):774. doi:10.3390/cells9030774 PubMed PMID: 32235738; PubMed Central PMCID: PMC7140691.

80. Urciuoli E, D’Oria V, Petrini S, Peruzzi B. Lamin A/C Mechanosensor Drives Tumor Cell Aggressiveness and Adhesion on Substrates With Tissue-Specific Elasticity. Front Cell Dev Biol. 2021;9:712377. doi:10.3389/fcell.2021.712377 PubMed PMID: 34595168; PubMed Central PMCID: PMC8476891.

81. Guelen L, Pagie L, Brasset E, Meuleman W, Faza MB, Talhout W, et al. Domain organization of human chromosomes revealed by mapping of nuclear lamina interactions. Nature. 2008 Jun 12;453(7197):948–51. doi:10.1038/nature06947 PubMed PMID: 18463634.

82. Kind J, Pagie L, de Vries SS, Nahidiazar L, Dey SS, Bienko M, et al. Genome-wide maps of nuclear lamina interactions in single human cells. Cell. 2015 Sep 24;163(1):134–47. doi:10.1016/j.cell.2015.08.040 PubMed PMID: 26365489; PubMed Central PMCID: PMC4583798.

83. Towbin BD, González-Aguilera C, Sack R, Gaidatzis D, Kalck V, Meister P, et al. Step-wise methylation of histone H3K9 positions heterochromatin at the nuclear periphery. Cell. 2012 Aug 31;150(5):934–47. doi:10.1016/j.cell.2012.06.051 PubMed PMID: 22939621.

84. Solovei I, Wang AS, Thanisch K, Schmidt CS, Krebs S, Zwerger M, et al. LBR and lamin A/C sequentially tether peripheral heterochromatin and inversely regulate differentiation. Cell. 2013 Jan 31;152(3):584–98. doi:10.1016/j.cell.2013.01.009 PubMed PMID: 23374351.

85. van Steensel B, Belmont AS. Lamina-Associated Domains: Links with Chromosome Architecture, Heterochromatin, and Gene Repression. Cell. 2017 May 18;169(5):780–91. doi:10.1016/j.cell.2017.04.022 PubMed PMID: 28525751; PubMed Central PMCID: PMC5532494.

86. Uhler C, Shivashankar GV. Chromosome Intermingling: Mechanical Hotspots for Genome Regulation. Trends Cell Biol. 2017 Nov;27(11):810–9. doi:10.1016/j.tcb.2017.06.005 PubMed PMID: 28728836.

87. Stephens AD, Banigan EJ, Adam SA, Goldman RD, Marko JF. Chromatin and lamin A determine two different mechanical response regimes of the cell nucleus. Mol Biol Cell. 2017 Jul 7;28(14):1984–96. doi:10.1091/mbc.E16-09-0653 PubMed PMID: 28057760; PubMed Central PMCID: PMC5541848.

88. Attar AG, Paturej J, Sarıyer OS, Banigan EJ, Erbaş A. Peripheral heterochromatin tethering is required for chromatin-based nuclear mechanical response. Nucleic Acids Res. 2025 Aug 11;53(15):gkaf763. doi:10.1093/nar/gkaf763 PubMed PMID: 40823810; PubMed Central PMCID: PMC12359041.

89. Hobson CM, Kern M, O’Brien ET, Stephens AD, Falvo MR, Superfine R. Correlating nuclear morphology and external force with combined atomic force microscopy and light sheet imaging separates roles of chromatin and lamin A/C in nuclear mechanics. Mol Biol Cell. 2020 Jul 21;31(16):1788–801. doi:10.1091/mbc.E20-01-0073 PubMed PMID: 32267206; PubMed Central PMCID: PMC7521857.

90. Xu J, Ma H, Jin J, Uttam S, Fu R, Huang Y, et al. Super-Resolution Imaging of Higher-Order Chromatin Structures at Different Epigenomic States in Single Mammalian Cells. Cell Rep. 2018 Jul 24;24(4):873–82. doi:10.1016/j.celrep.2018.06.085 PubMed PMID: 30044984; PubMed Central PMCID: PMC6154382.

91. Shah P, Hobson CM, Cheng S, Colville MJ, Paszek MJ, Superfine R, et al. Nuclear Deformation Causes DNA Damage by Increasing Replication Stress. Curr Biol. 2021 Feb 22;31(4):753–765.e6. doi:10.1016/j.cub.2020.11.037 PubMed PMID: 33326770; PubMed Central PMCID: PMC7904640.

92. Fortuny A, Chansard A, Caron P, Chevallier O, Leroy O, Renaud O, et al. Imaging the response to DNA damage in heterochromatin domains reveals core principles of heterochromatin maintenance. Nat Commun. 2021 Apr 23;12(1):2428. doi:10.1038/s41467-021-22575-5 PubMed PMID: 33893291; PubMed Central PMCID: PMC8065061.

93. Svobodová Kovaříková A, Legartová S, Krejčí J, Bártová E. H3K9me3 and H4K20me3 represent the epigenetic landscape for 53BP1 binding to DNA lesions. Aging (Albany NY). 2018 Oct 11;10(10):2585–605. doi:10.18632/aging.101572 PubMed PMID: 30312172; PubMed Central PMCID: PMC6224238.

94. Sun Y, Jiang X, Xu Y, Ayrapetov MK, Moreau LA, Whetstine JR, et al. Histone H3 methylation links DNA damage detection to activation of the tumour suppressor Tip60. Nat Cell Biol. 2009 Nov;11(11):1376–82. doi:10.1038/ncb1982 PubMed PMID: 19783983; PubMed Central PMCID: PMC2783526.

95. Ayrapetov MK, Gursoy-Yuzugullu O, Xu C, Xu Y, Price BD. DNA double-strand breaks promote methylation of histone H3 on lysine 9 and transient formation of repressive chromatin. Proc Natl Acad Sci U S A. 2014 Jun 24;111(25):9169–74. doi:10.1073/pnas.1403565111 PubMed PMID: 24927542; PubMed Central PMCID: PMC4078803.

96. Elosegui-Artola A, Andreu I, Beedle AEM, Lezamiz A, Uroz M, Kosmalska AJ, et al. Force Triggers YAP Nuclear Entry by Regulating Transport across Nuclear Pores. Cell. 2017 Nov 30;171(6):1397–1410.e14. doi:10.1016/j.cell.2017.10.008 PubMed PMID: 29107331.

97. Ghagre A, Delarue A, Srivastava LK, Koushki N, Ehrlicher A. Nuclear curvature determines Yes-associated protein localization and differentiation of mesenchymal stem cells. Biophys J. 2024 May 21;123(10):1222–39. doi:10.1016/j.bpj.2024.04.008 PubMed PMID: 38605521; PubMed Central PMCID: PMC11140468.

98. Chen J, Tsai YH, Linden AK, Kessler JA, Peng CY. YAP and TAZ differentially regulate postnatal cortical progenitor proliferation and astrocyte differentiation. Journal of Cell Science. 2024 May 15;137(10):jcs261516. doi:10.1242/jcs.261516

99. Sun C, De Mello V, Mohamed A, Ortuste Quiroga HP, Garcia-Munoz A, Al Bloshi A, et al. Common and Distinctive Functions of the Hippo Effectors Taz and Yap in Skeletal Muscle Stem Cell Function. Stem Cells. 2017 Aug 1;35(8):1958–72. doi:10.1002/stem.2652

100. Koushki N, Ghagre A, Srivastava LK, Molter C, Ehrlicher AJ. Nuclear compression regulates YAP spatiotemporal fluctuations in living cells. Proc Natl Acad Sci U S A. 2023 Jul 11;120(28):e2301285120. doi:10.1073/pnas.2301285120 PubMed PMID: 37399392; PubMed Central PMCID: PMC10334804.

101. Das A, Fischer RS, Pan D, Waterman CM. YAP Nuclear Localization in the Absence of Cell-Cell Contact Is Mediated by a Filamentous Actin-dependent, Myosin II- and Phospho-YAP-independent Pathway during Extracellular Matrix Mechanosensing. J Biol Chem. 2016 Mar 18;291(12):6096–110. doi:10.1074/jbc.M115.708313 PubMed PMID: 26757814; PubMed Central PMCID: PMC4813550.

102. Jain N, Iyer KV, Kumar A, Shivashankar GV. Cell geometric constraints induce modular gene-expression patterns via redistribution of HDAC3 regulated by actomyosin contractility. Proc Natl Acad Sci USA. 2013 Jul 9;110(28):11349–54. doi:10.1073/pnas.1300801110

103. Li Q, Verma IM. NF-kappaB regulation in the immune system. Nat Rev Immunol. 2002 Oct;2(10):725–34. doi:10.1038/nri910 PubMed PMID: 12360211.

104. Xue J, Yang XR, Wang L. NF-κB signaling pathway in osteosarcoma: from signaling networks to targeted therapy. Front Oncol. 2025;15:1565760. doi:10.3389/fonc.2025.1565760 PubMed PMID: 40786506; PubMed Central PMCID: PMC12333507.

105. Zeman MK, Cimprich KA. Causes and consequences of replication stress. Nat Cell Biol. 2014 Jan;16(1):2–9. doi:10.1038/ncb2897 PubMed PMID: 24366029; PubMed Central PMCID: PMC4354890.

106. Zhao J, Zhang L, Lu A, Han Y, Colangelo D, Bukata C, et al. ATM is a key driver of NF-κB-dependent DNA-damage-induced senescence, stem cell dysfunction and aging. Aging (Albany NY). 2020 Mar 22;12(6):4688–710. doi:10.18632/aging.102863 PubMed PMID: 32201398; PubMed Central PMCID: PMC7138542.

107. McCool KW, Miyamoto S. DNA damage-dependent NF-κB activation: NEMO turns nuclear signaling inside out. Immunol Rev. 2012 Mar;246(1):311–26. doi:10.1111/j.1600-065X.2012.01101.x PubMed PMID: 22435563; PubMed Central PMCID: PMC3311051.

108. Podhorecka M, Skladanowski A, Bozko P. H2AX Phosphorylation: Its Role in DNA Damage Response and Cancer Therapy. J Nucleic Acids. 2010 Aug 3;2010:920161. doi:10.4061/2010/920161 PubMed PMID: 20811597; PubMed Central PMCID: PMC2929501.

109. Marcellus RC, Teodoro JG, Charbonneau R, Shore GC, Branton PE. Expression of p53 in Saos-2 osteosarcoma cells induces apoptosis which can be inhibited by Bcl-2 or the adenovirus E1B-55 kDa protein. Cell Growth Differ. 1996 Dec;7(12):1643–50. PubMed PMID: 8959332.

110. Lukin DJ, Carvajal LA, Liu W jun, Resnick-Silverman L, Manfredi JJ. p53 Promotes cell survival due to the reversibility of its cell-cycle checkpoints. Mol Cancer Res. 2015 Jan;13(1):16–28. doi:10.1158/1541-7786.MCR-14-0177 PubMed PMID: 25158956; PubMed Central PMCID: PMC4312522.

111. Bartkova J, Horejsí Z, Koed K, Krämer A, Tort F, Zieger K, et al. DNA damage response as a candidate anti-cancer barrier in early human tumorigenesis. Nature. 2005 Apr 14;434(7035):864–70. doi:10.1038/nature03482 PubMed PMID: 15829956.

112. Halazonetis TD, Gorgoulis VG, Bartek J. An oncogene-induced DNA damage model for cancer development. Science. 2008 Mar 7;319(5868):1352–5. doi:10.1126/science.1140735 PubMed PMID: 18323444.

113. Luu AK, Viloria-Petit AM. Targeting Mechanotransduction in Osteosarcoma: A Comparative Oncology Perspective. Int J Mol Sci. 2020 Oct 14;21(20):7595. doi:10.3390/ijms21207595 PubMed PMID: 33066583; PubMed Central PMCID: PMC7589883.

114. Soni L, Kaur A, Sharma A. A Review on Different Versions and Interfaces of Blender Software. In: 2023 7th International Conference on Trends in Electronics and Informatics (ICOEI) [Internet]. Tirunelveli, India: IEEE; 2023 [cited 2026 Feb 23]. p. 882–7. Available from: https://ieeexplore.ieee.org/document/10125672/ doi:10.1109/ICOEI56765.2023.10125672

115. De Feraudy S, Revet I, Bezrookove V, Feeney L, Cleaver JE. A minority of foci or pan-nuclear apoptotic staining of γH2AX in the S phase after UV damage contain DNA double-strand breaks. Proc Natl Acad Sci USA. 2010 Apr 13;107(15):6870–5. doi:10.1073/pnas.1002175107

116. Kwon JW, Kwon HK, Shin HJ, Choi YM, Anwar MA, Choi S. Activating transcription factor 3 represses inflammatory responses by binding to the p65 subunit of NF-κB. Sci Rep. 2015 Sep 28;5(1):14470. doi:10.1038/srep14470

117. Yurube T, Buchser WJ, Moon HJ, Hartman RA, Takayama K, Kawakami Y, et al. Serum and nutrient deprivation increase autophagic flux in intervertebral disc annulus fibrosus cells: an in vitro experimental study. Eur Spine J. 2019 May;28(5):993–1004. doi:10.1007/s00586-019-05910-9 PubMed PMID: 30847707; PubMed Central PMCID: PMC6538458.

118. Ghali M, Brahmi C, Benltifa M, Vaulot C, Airoudj A, Fioux P, et al. Characterization of polyoxometalate/polymer photo-composites: A toolbox for the photodegradation of organic pollutants. Journal of Polymer Science. 2021 Jan 15;59(2):153–69. doi:10.1002/pol.20200568

119. Sader JE, Larson I, Mulvaney P, White LR. Method for the calibration of atomic force microscope cantilevers. Review of Scientific Instruments. 1995 Jul 1;66(7):3789–98. doi:10.1063/1.1145439

120. Noursadeghi M, Tsang J, Haustein T, Miller RF, Chain BM, Katz DR. Quantitative imaging assay for NF-kappaB nuclear translocation in primary human macrophages. J Immunol Methods. 2008 Jan 1;329(1–2):194–200. doi:10.1016/j.jim.2007.10.015 PubMed PMID: 18036607; PubMed Central PMCID: PMC2225449.

121. Lapytsko A, Kollarovic G, Ivanova L, Studencka M, Schaber J. FoCo: a simple and robust quantification algorithm of nuclear foci. BMC Bioinformatics. 2015 Nov 21;16:392. doi:10.1186/s12859-015-0816-5 PubMed PMID: 26589438; PubMed Central PMCID: PMC4654864.

122. Memmel S, Sisario D, Zimmermann H, Sauer M, Sukhorukov VL, Djuzenova CS, et al. FocAn: automated 3D analysis of DNA repair foci in image stacks acquired by confocal fluorescence microscopy. BMC Bioinformatics. 2020 Jan 28;21(1):27. doi:10.1186/s12859-020-3370-8 PubMed PMID: 31992200; PubMed Central PMCID: PMC6986076.

123. Vassaux M, Milan JL. Stem cell mechanical behaviour modelling: substrate’s curvature influence during adhesion. Biomech Model Mechanobiol. 2017 Aug;16(4):1295–308. doi:10.1007/s10237-017-0888-4

124. Vassaux M. mvassaux/adhSC: Initial release of the adhesion cell model [Internet]. Zenodo; 2018 [cited 2026 Feb 23]. Available from: https://zenodo.org/record/1187087 doi:10.5281/ZENODO.1187087

125. Dubois F, Jean M. The non smooth contact dynamic method: recent LMGC90 software developments and application. In: Wriggers P, Nackenhorst U, editors. Analysis and Simulation of Contact Problems [Internet]. Berlin/Heidelberg: Springer-Verlag; 2006 [cited 2026 Feb 23]. p. 375–8. (Lecture Notes in Applied and Computational Mechanics). Available from: http://link.springer.com/10.1007/3-540-31761-9_44 doi:10.1007/3-540-31761-9_44

126. Lherbette M, Dos Santos Á, Hari-Gupta Y, Fili N, Toseland CP, Schaap IAT. Atomic Force Microscopy micro-rheology reveals large structural inhomogeneities in single cell-nuclei. Sci Rep. 2017 Aug 14;7(1):8116. doi:10.1038/s41598-017-08517-6 PubMed PMID: 28808261; PubMed Central PMCID: PMC5556037.

