## Supplemental Data 1 for "Nucleus confinement within concave microcavities modulates nuclear morphology, subnuclear dynamics and mechanotransduction in human osteosarcoma cells"

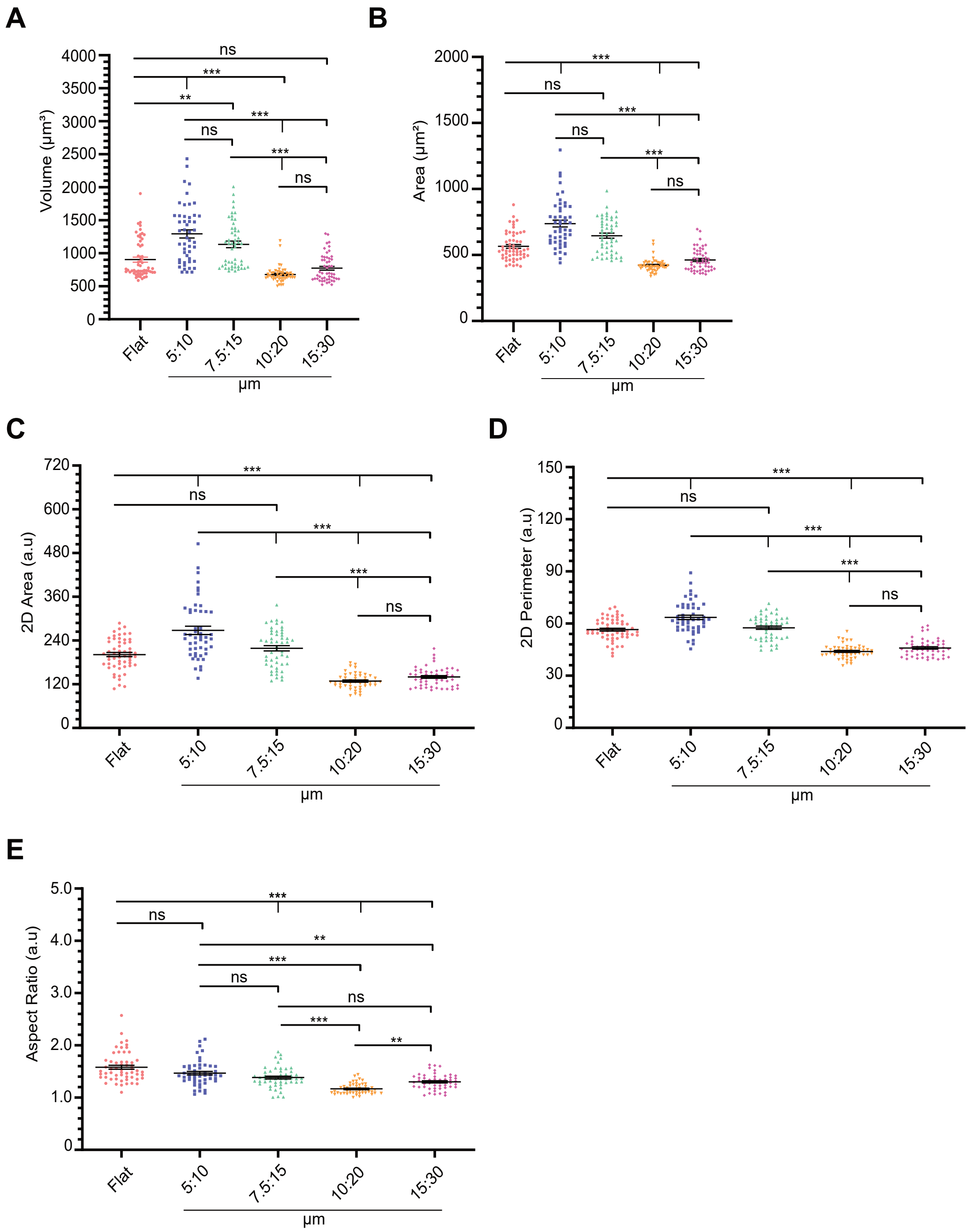
A
Volume (µm³)
ns
***
**
ns
***
***
ns
B
Area (µm²)
***
ns
***
ns
***
ns
C
2D Area (a.u)
***
ns
***
***
ns
D
2D Perimeter (a.u)
***
ns
***
***
ns
E
Aspect Ratio (a.u)
***
ns
**
***
ns
ns
***
**
Flat
5:10
7.5:15
10:20
15:30
µm

SI. Figure S1

A

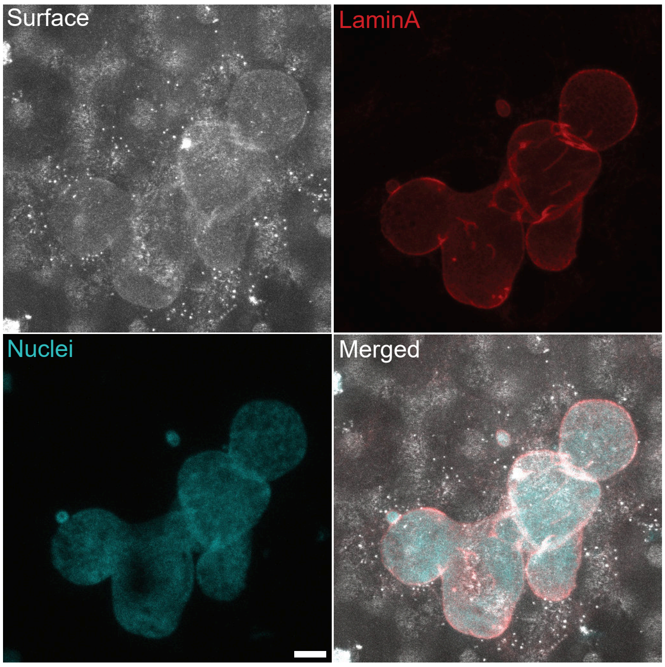


B

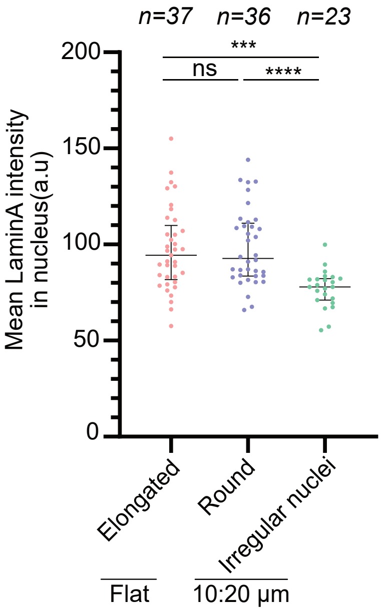


C

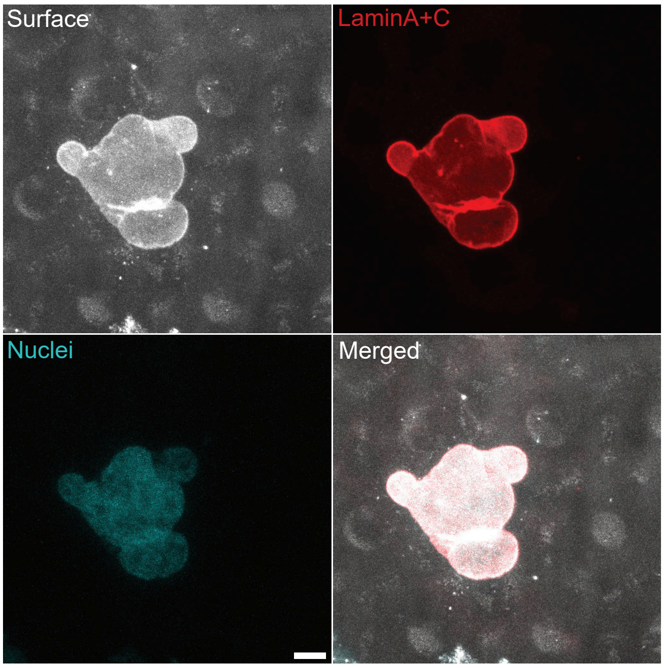


D

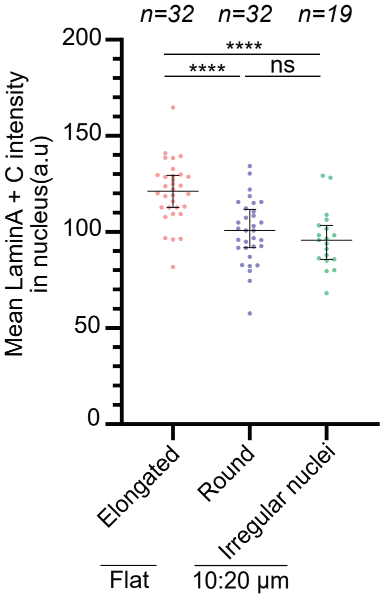


SI Figure S2

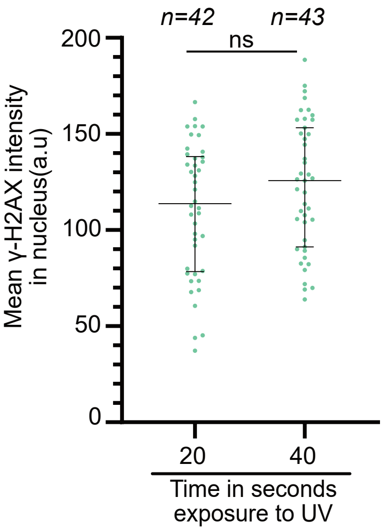
n=42
n=43
ns
200
150
100
50
0
Mean γ-H2AX intensity
in nucleus(a.u)
20
40
Time in seconds
exposure to UV

SI Figure S3

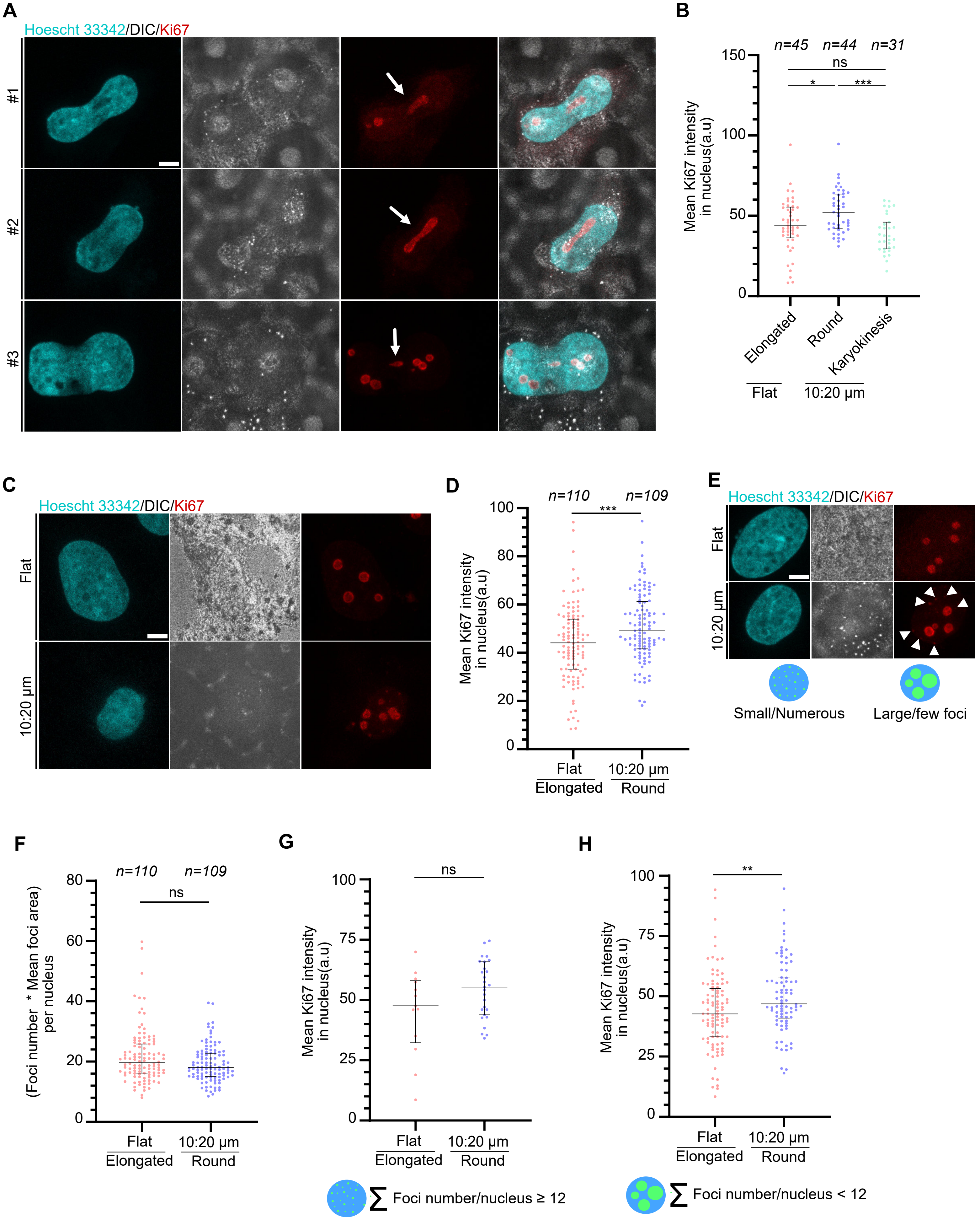
A
Hoescht 33342/DIC/Ki67
#1
#2
#3
B
n=45
n=44
n=31
ns
*
***
150
100
50
0
Mean Ki67 intensity
in nucleus(a.u)
Elongated
Round
Karyokinesis
Flat
10:20 µm
C
Hoescht 33342/DIC/Ki67
Flat
10:20 µm
D
n=110
n=109
***
100
80
60
40
20
0
Mean Ki67 intensity
in nucleus(a.u)
Flat
Elongated
10:20 µm
Round
E
Hoescht 33342/DIC/Ki67
Flat
10:20 µm
Small/Numerous
Large/few foci
F
n=110
n=109
ns
80
60
40
20
0
(Foci number * Mean foci area)
per nucleus
Flat
Elongated
10:20 µm
Round
G
ns
100
75
50
25
0
Mean Ki67 intensity
in nucleus(a.u)
Flat
Elongated
10:20 µm
Round
∑ Foci number/nucleus ≥ 12
H
**
100
75
50
25
0
Mean Ki67 intensity
in nucleus(a.u)
Flat
Elongated
10:20 µm
Round
∑ Foci number/nucleus < 12

SI Figure S4

| Sample | Elasticity (GPa-MPa) |
|---|---|
| Polystrene | 0.53751 ± 0.0167 GPa |
| PDMS (1:10) | 5.26955 ± 4.6723 MPa |
| PDMS (1:50) | 0.2377 ± 0.01 MPa |

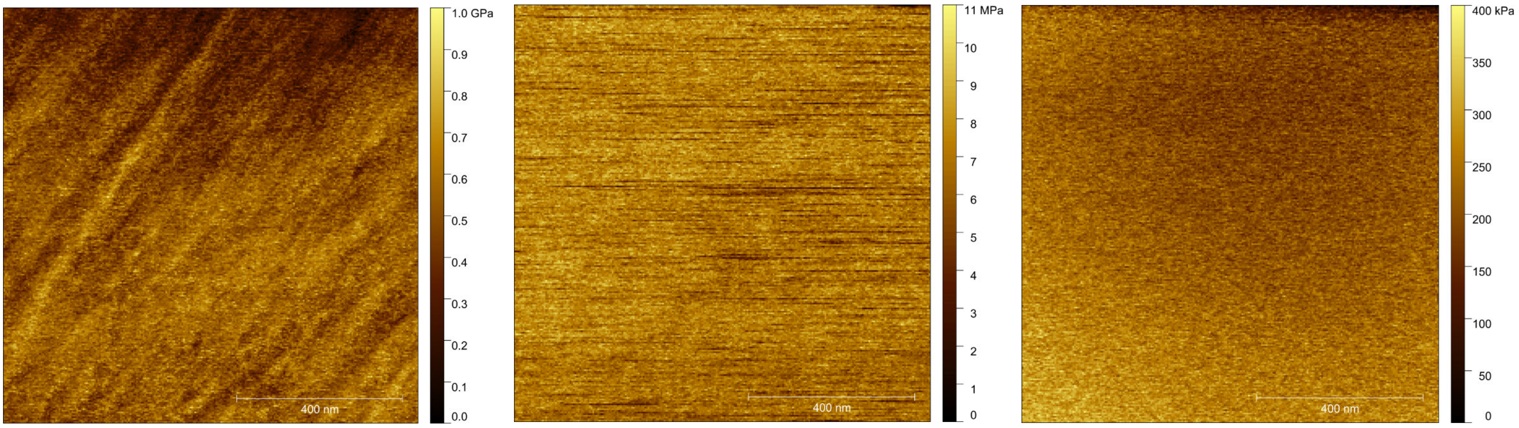


Polystrene PDMS (1:10) PDMS (1:50)

SI Figure S5

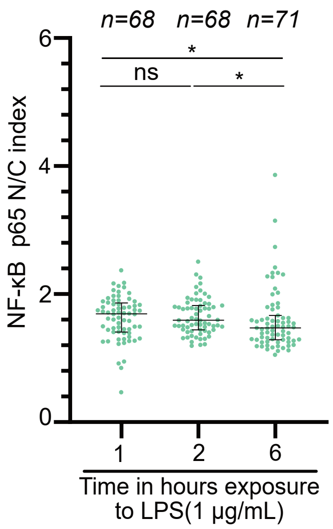
n=68
n=68
n=71
*
ns
*
NF-κB p65 N/C index
6
4
2
0
1
2
6
Time in hours exposure
to LPS(1 μg/mL)

SI Figure S6
